# Cryo-EM structure of the *Arabidopsis thaliana* V-type ATPase

**DOI:** 10.64898/2026.07.01.735876

**Authors:** Mariia Khamina, Nadja Wunsch, Upendo Lupanga, Fabian Fink, Hanlin Wang, Waltraud X. Schulze, Karin Schumacher, John L. Rubinstein

## Abstract

Vacuolar-type ATPases (V-ATPases) are evolutionarily conserved rotary proton pumps that play essential roles in the eukaryotic cell. By coupling ATP hydrolysis in their cytosolic V_1_ region to proton translocation through their membrane-embedded V_O_ region, V-ATPases establish and maintain an acidic pH in the lumen of several different organelles. Functional diversity in the pump is enabled by multiple paralogous genes for the subunits of the complex, which are expressed in a tissue- and organelle-specific manner. Interactions between V-ATPase and TLDc domain-containing proteins have been shown to regulate the enzyme in yeast and mammals but their relevance in plants has remained unclear. We isolated the endogenous V-ATPase from *Arabidopsis thaliana* leaves and determined its structure by electron cryomicroscopy. Mass spectrometry showed that most of the enzyme originated from the tonoplast. The structural analysis revealed the full rotary catalytic cycle of the plant V-ATPase, and a combination of structural and biochemical experiments showed S-acylation of subunits AP1 and the tonoplast-specific subunit a3 isoform. A subpopulation of complexes derived from the trans-Golgi network/early endosome was identified and found to bind the TLDc protein OXR5. Together, these findings reveal plant-specific features in V-ATPase and suggest organelle-specific interactions with TLDc proteins, pointing to conserved but context-dependent V-ATPase regulation in eukaryotes.

## Introduction

Within organelles in eukaryotic cells, pH is a tightly controlled biophysical parameter that helps define the compartment’s role in cellular homeostasis. The neutral pH within the endoplasmic reticulum, and the pH gradient across the Golgi allow protein maturation and trafficking. The plant *trans* Golgi network/early endosome (TGN/EE), a distinct organelle that receives and sorts proteins from the endocytic, recycling, and secretory pathways, is also maintained at an acidic pH (Viotti et al., 2010). Acidified vacuoles process and store nutrients and help degrade pathogens. Plants also use these acidic vacuoles to adapt to environmental stressors such as salinity, drought, heavy metals, and extreme temperatures (Apse et al., 1999; Catalá et al., 2003; Park et al., 2012; Schulze et al., 2012; Sun et al., 2025; Li et al., 2022; Francis et al., 2005). Tolerance to these abiotic stressors is supported by the electrochemical proton gradient across the vacuolar membrane, known as the tonoplast (Martinoia et al., 2007). This proton motive force energizes secondary transporters responsible for sequestering ions and toxic compounds and drives osmotic water influx into the vacuole, providing turgor pressure within the cell. In fact, vacuoles can occupy up to 90% of cell volume, enabling plants to grow to large sizes through water uptake rather than synthesis of more metabolically expensive cytosol (Krüger and Schumacher, 2018).

Vacuolar-type ATPases (V-ATPases), along with vacuolar pyrophosphatases, are proton pumps that contribute to acidification of plant organelles (Kriegel et al., 2015). V-ATPases are multiprotein complexes comprising a soluble V_1_ region (subunits VHA-A, -B, -C, -D, -E, -F, -G, and -H, hereafter referred to without the VHA prefix) and a membrane-embedded V_O_ region (in plants, subunits VHA-a, -c, -c″, -d, -e, -AP1, and -AP2). Three pairs of catalytic A and B subunits sequentially hydrolyze ATP molecules causing rotation of a central rotor subcomplex, which extends from the V_1_ region (subunits D and F) into the V_O_ region (subunits d and a c-ring, formed from nine c subunits and subunit c″). Rotation of the c-ring against subunit a in the V_O_ region drives proton translocation from the cytosol into the lumen of the organelle. Three peripheral stalk structures, each formed from subunits E and G, hold subunit a stationary relative to the A_3_B_3_ hexamer. Many of the subunits that are not involved directly in ATP hydrolysis or rotation-driven proton translocation participate in stabilization and regulation of the protein complex (Hong-Hermesdorf et al., 2006; Vasanthakumar et al., 2022; Chen et al., 2026).

Several plant V-ATPase subunits have multiple isoforms, encoded by gene paralogs (Sze et al., 2002). The number of isoforms for each subunit varies between plant species and physiological functions for many of the isoforms are not fully understood (Schumacher and Krebs, 2010). In the model plant *Arabidopsis thaliana*, different isoforms of subunits G and E are expressed in different tissues (Strompen et al., 2005; Hanitzsch et al., 2007; Dettmer et al., 2010). In contrast, isoforms of subunits B and d are found in the same tissue but are expressed differentially in the presence of environmental stressors (Hanitzsch et al., 2007; Feng et al., 2020). Isoforms of subunit a are found in distinct organelles: subunit a1 localizes to the TGN/EE, while subunits a2 and a3 are both found in tonoplasts (Dettmer et al., 2006; Krebs et al., 2010; Lupanga et al., 2020). This variability in V-ATPase subunit isoforms likely provides spatial and temporal control of the proton pump.

A major mechanism of V-ATPase regulation is the reversible dissociation of the enzyme’s V_1_ region from its V_O_ region, with ATP hydrolysis in the isolated V_1_ complex inhibited and the V_O_ complex becoming impermeable to protons (Kane and Parra, 2000; Qi and Forgac, 2008). In the yeast *Saccharomyces cerevisiae*, glucose depletion in the growth medium leads to reversible dissociation of V_1_ from V_O_ (Kane, 1995). However, in plants, neither reversible dissociation of the V-ATPase nor the environmental signals that induce this process have been identified. V-ATPase activity can also be regulated by proteins that contain Tre2/Bub2/Cdc16 lysin motif domain catalytic (TLDc)-domains. TLDc proteins were first identified as protective against oxidative stress (Volkert et al., 2000) but more recently have been recognized as conserved interactors of V-ATPases in insects, mammals, and yeast (Merkulova et al., 2015; Eaton et al., 2021). While the precise functions of TLDc proteins are still unclear, they have been shown to affect the assembly status of the proton pump (Khan et al., 2025, 2022; Klössel et al., 2024; Oot and Wilkens, 2024). The Arabidopsis genome encodes six TLDc proteins, designated OXR1 to OXR6 (Colombatti et al., 2019). Like TLDc proteins in other organisms, plant OXR2 was described as protective against oxidative and biotic stress (Colombatti et al., 2019; Mencia et al., 2020). However, to our knowledge, the interaction of plant TLDc proteins with V-ATPases have not been investigated.

Previous structural studies of V-ATPases from mammals and yeast not only revealed the proton-pumping mechanism of the complex, but also new interacting partners (Abbas et al., 2020; Coupland et al., 2024; Mazhab-Jafari et al., 2016; Roh et al., 2018; Tan et al., 2022a; Vasanthakumar et al., 2022; L. Wang et al., 2022; R. Wang et al., 2022; Zhao et al., 2015). A first structure of a plant V-ATPase was determined from citrus fruit (Tan et al., 2022b), defining the overall architecture of the enzyme. The citrus V-ATPase has a V_O_ subunit composition distinct from the mammalian and yeast proton pumps. In addition, the citrus V-ATPase structure revealed a unique conformation of the H subunit in the V_1_ region, suggesting an additional level of activity regulation. However, resolution of that structure was insufficient to show the full rotary cycle, prominent posttranslational modifications, and did not show any interacting proteins.

We isolated the V-ATPase from Arabidopsis and used cryo-EM to determine its structure. Overall, the structure resembles the earlier structure from citrus fruit, but at higher resolution and with improved map density for the subunits of the complex. The structure, along with biochemical analysis, revealed plant-specific post-translational modifications, with the AP1 subunit and tonoplast a3 subunit acylated at the sulfur atoms of several cysteine residues. Further, endogenous OXR5 was bound to a subpopulation of complexes, inducing a strained conformation of V-ATPase that may modulate its activity. We demonstrate that OXR5 is only associated with the V-ATPase at the TGN/EE in Arabidopsis, suggesting that V-ATPases in different subcellular locations are subjected to differential modulation by TLDc proteins.

## Results

### Structure of Arabidopsis V-ATPase reveals the enzyme in three rotational states

Cellular membranes from Arabidopsis rosette leaves were collected and solubilized with detergent. V-ATPase was isolated from these membranes using recombinant SidK, a *Legionella pneumophila* effector protein used previously to isolate mammalian (Abbas et al., 2020), yeast (Maxson et al., 2022), and plant (Tan et al., 2022b) V-ATPase. The purified protein (**Fig. 1A**) was subjected to structural analysis by cryo-EM, which yielded multiple maps of the enzyme. The best of these maps was at 3.0 Å resolution, with focused refinement of the V_1_, V_O_, and peripheral stalk regions of the complex reaching resolutions of 2.7, 2.8, and 3.6 Å, respectively (**Fig. 1B**, **Fig. S1**, **Fig. S2**, and **Table S1**). The structure showed that Arabidopsi*s* V-ATPase has an architecture similar to orthologous proton pumps in mammals, yeast, and citrus fruit (Wang and Rubinstein, 2023). Like the citrus fruit V-ATPase structure, density for SidK is not seen in the cryo-EM map of Arabidopsis V-ATPase, indicating that SidK binds the plant enzyme with lower affinity than the yeast and mammalian enzymes.

**Figure 1.**
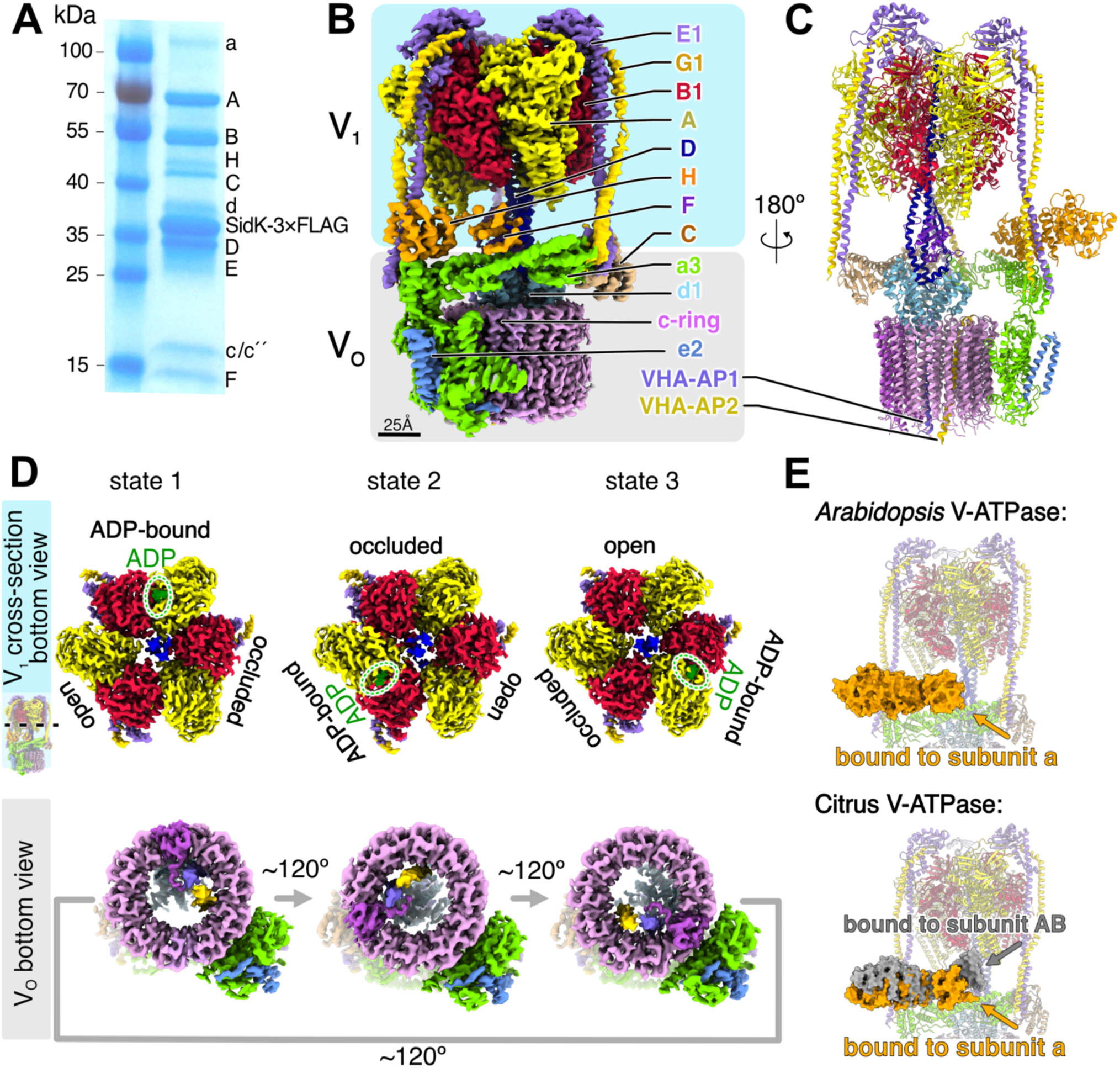
Overall structure of *Arabidopsis* V-ATPase. (A) SDS-PAGE of purified V-ATPase. (B) Composite map of Arabidopsis V-ATPase in rotational state 1. (C) Atomic model of Arabidopsis V-ATPase in rotational state 1. (D) Cross sections of the V_1_ and V_O_ regions in three rotational states. (E) Conformations of subunit H in V-ATPase structures of Arabidopsis V-ATPase (*top*) and the citrus fruit (*bottom*) V-ATPase.

The cryo-EM structure, combined with mass spectrometry of the protein preparation (**Supplementary Data 1**), identified the subunit composition of the protein complex as A_3_B1_3_CDE1_3_FG1_3_H in the V_1_ region, and subunits a3, c_9_, c″, d1, AP1, and AP2 in the V_O_ region. Subunit e was not identified by mass spectrometry but was modelled as isoform e2 based on sidechain densities observed in the cryo-EM map (**Fig. S3A**). The presence of the a3 isoform indicates that the predominant V-ATPase species isolated was from the tonoplast (Carter et al., 2004; Jaquinod et al., 2007). The presence of the E1 and G1 isoforms is consistent with the protein complex being isolated from the leaves of Arabidopsis plants, where these isoforms are highly expressed (Strompen et al., 2005; Hanitzsch et al., 2007; Dettmer et al., 2010). The subunit isoforms B2, E3, G2, a1, a2, and d2 were also detected by mass spectrometry of the sample, but their low iBAQ scores (Schwanhäusser et al., 2011; Zauber et al., 2013) suggest that they are minor components of the preparation.

The resolution of the cryo-EM map, combined with information about the subunit isoforms from mass spectrometry, was sufficient to build an atomic model comprising 95 % of the amino acid residues of the complex (**Fig. 1C** and **Table S2**). Cryo-EM of yeast and mammalian V-ATPases has shown that, in solution without free ATP, these enzymes adopt three main rotational states, known as rotational state 1, state 2, and state 3 (Abbas et al., 2020; Zhao et al., 2015). However, structural studies of the citrus V-ATPase (Tan et al., 2022b) were only able to identify two of the three states. The Arabidopsis enzyme imaged here was found in all three rotational states (**Fig. 1D**). Together, the three resulting atomic models describe the full rotational cycle of the plant V-ATPase. In this cycle (Abbas et al., 2020; Zhao et al., 2015), one of the three catalytic pairs of subunits A and B adopts an “open” conformation that binds and hydrolyses an ATP molecule to ADP. ATP hydrolysis causes a ∼120° rotation (clockwise when viewed from V_1_ toward V_O_) of the central rotor (subunits D, F, d, and the c ring) with the AB pair subsequently adopting an “ADP-bound” conformation. One of the two adjacent AB pairs, which was initially in the ADP-bound conformation, releases its nucleotide and takes on an “occluded” conformation.

Simultaneously, the third AB pair, which was in the occluded conformation, transitions to the open conformation primed for the next step in the catalytic cycle. Density corresponding to ATP is not present in the Arabidopsis V-ATPase cryo-EM map, consistent with the absence of free ATP in solution during sample preparation. However, density corresponding to ADP, the product of the catalytic cycle, is apparent in one AB pair in each of the three rotational states (**Fig. 1D**, *top row*, *green ovals*).

Subunit H plays an essential regulatory role during reversible dissociation of V-ATPase, inhibiting ATP hydrolysis by the isolated V_1_ complex (Parra et al., 2000). This V_1_ inhibition is accomplished by subunit H bridging peripheral stalks #1 and #2 in the isolated V_1_ complex, which prevents the conformational changes needed for ATP hydrolysis (Vasanthakumar et al., 2022) (PDB ID 7UZJ). In the assembled and ATP-hydrolysis-competent V-ATPases from yeast and mammals, the N-terminal domain of subunit H binds peripheral stalk #1 as well as subunit a from the V_O_ region while its C-terminal domain only binds subunit a (Benlekbir et al., 2012; Wang et al., 2020). In contrast, in the intact citrus fruit V-ATPase, subunit H is more dynamic, with the C-terminal domain of subunit H able to bind either to subunit a (**Fig. 1E**, *lower*, *orange conformation*) or to subunits A and B from the V_1_ region (**Fig. 1E**, *lower*, *grey conformation*) (Tan et al., 2022b). This second conformation of the citrus V-ATPase could not be detected for the Arabidopsis V-ATPase, which behaves more like the proton pump in mammals and yeast and only binds subunit a (**Fig. 1E**, *upper*, *orange conformation*).

### The Arabidopsis V-ATPase V_O_ region has multiple post-translational modifications

The subunit composition of the V-ATPase V_O_ region is not completely conserved across eukaryotes. Specifically, the small transmembrane subunits from the complex differ between kingdoms (Abbas et al., 2020; Mazhab-Jafari et al., 2016; Roh et al., 2018; Tan et al., 2022b). The membrane-embedded c-ring in yeast has eight c subunits, subunit c′, and subunit c″, while plants and mammals have nine c subunits and subunit c″. Within the c ring, mammalian V-ATPase has transmembrane α helices from subunit c″, ATP6AP2/PRR, and ATP6AP1/Ac45 (Abbas et al., 2020), the latter also possessing a large luminal domain, while yeast V-ATPase has only transmembrane α helice from subunit c″ and Voa1p (Mazhab-Jafari et al., 2016; Roh et al., 2018). Like the enzyme from citrus fruit, Arabidopsis V-ATPase lacks the membrane-embedded subunit f, which is present both in yeast and mammals. The arrangement of subunits trapped within the Arabidopsis c-ring also resembles that of citrus V-ATPase, with well-resolved densities for subunits AP1 and AP2, and the absence of the N-terminal α helix from subunit c″ that is found in yeast and mammals (**Fig. 2A**).

**Figure 2.**
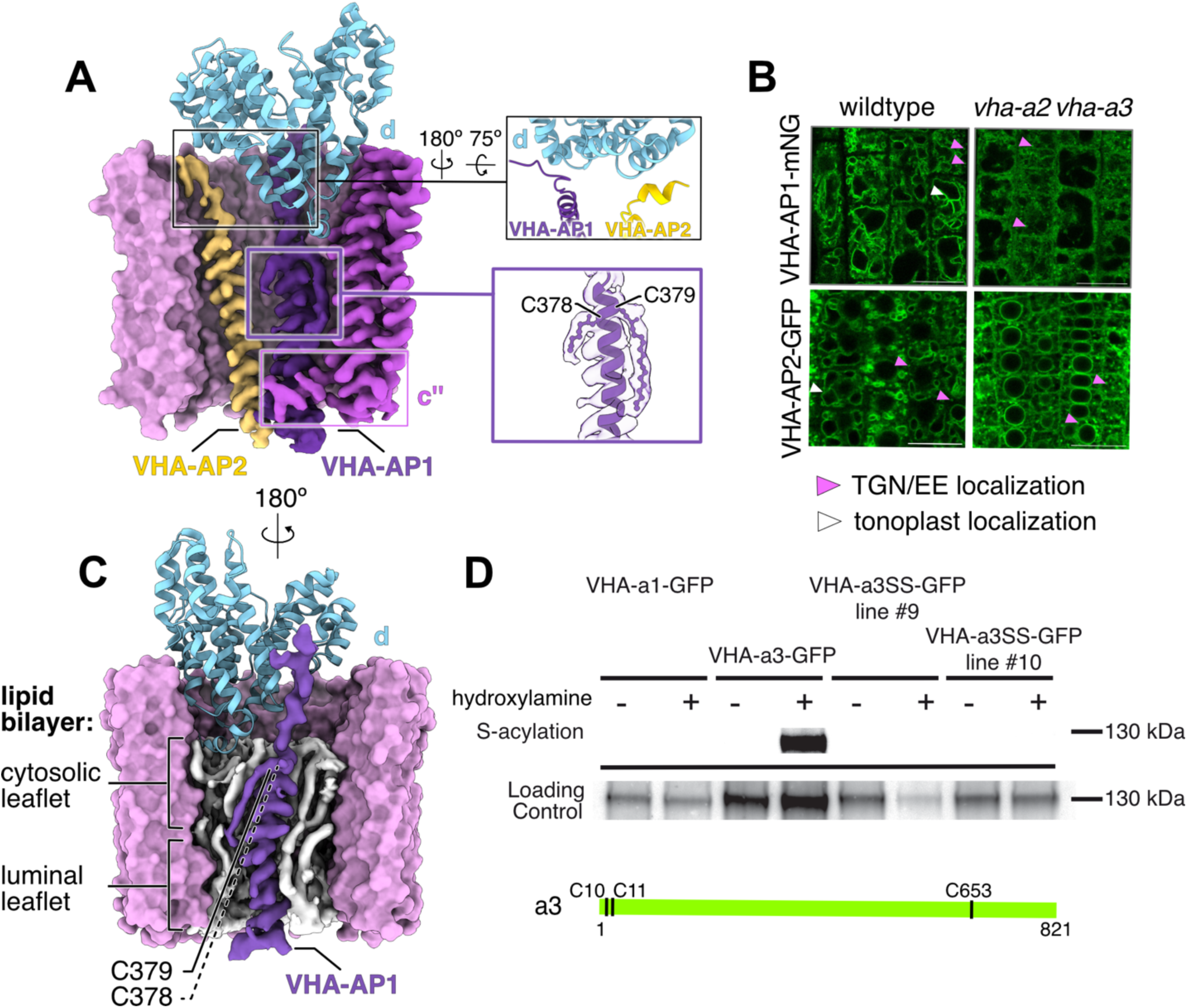
Architecture and post-translational modifications of the V_O_ subcomplex. (A) Structure of the c-ring (*top right*) showing interactions of AP1, AP2, and d subunits, and S-acylation of Cys378 and Cys379 in AP1 (*bottom right*). (B) In wild-type plants (*left*), AP1 and AP2 localize both to the vacuolar membrane and to small punctate structures, which are likely TGN/EE. In contrast, in the *vha-a2 vha-a3* mutant (*right*), where V-ATPases are absent from the vacuolar membrane, AP1 and AP2 are no longer detected at the tonoplast. Scale bar, 20 µm. (C) S-acylation modification of AP1 aligns with cytosolic leaflet of lipid bilayer inside the c-ring. (D) Domain structure and S-acylation sites of subunits AP1 and a3. (E) Western blot to detect S-acylation in subunit a1 (VHA-a1-GFP), subunit a3 (VHA-a3-GFP), and subunit a3 with Cys10 and Cys11 mutated to Ser (VHA-a3SS-GFP line #9 and VHA-a3SS-GFP line #10). All samples contain subunit a-GFP, as detected with an anti-GFP antibody (*upper*). After substituting S-acylation with biotin and purifying with neutravidin, biotinylation is detected only for VHA-a3-GFP (*bottom*).

The location of AP1 and AP2 within the lumen of the c-ring suggests that they are incorporated into the V_O_ complex during its initial biosynthesis (Abbas et al., 2020). In the Arabidopsis V-ATPase, AP1 and AP2 interact with subunit d1 (**Fig. 2A**, *top right*), which was not observed in the citrus V-ATPase structure. Orthologs of AP1 and AP2 in both yeast and mammals were hypothesized to facilitate assembly of V_O_ by anchoring the cytosolic subunit d to the c-ring (Kinouchi et al., 2010; Roh et al., 2018; Wang et al., 2023). We examined the intracellular distribution of AP1 and AP2 with confocal laser scanning microscopy using mNeonGreen (mNG)- and green fluorescent protein (GFP)-tagged variants of the proteins expressed under the ubiquitin-10 promoter (AP1-mNG, AP2-GFP). In wild-type plants, AP1-mNG and AP2-GFP localize both to the tonoplast and to small punctate structures, which are likely TGN/EE (**Fig. 2B**, *left*). In contrast, in a *vha-a2 vha-a3* mutant, where V-ATPases are absent from the tonoplast, AP1 and AP2 are no longer detected at the tonoplast, suggesting that their localization depends on the presence of V-ATPases (**Fig. 2B**, *right*) (Krebs et al., 2010).

Inspection of the cryo-EM map region corresponding to AP1 revealed elongated densities extending from residues Cys378 and Cys379, corresponding to S-acylation of the cysteines (**Fig. 2A**, *bottom right*). The density extending from Cys379 could be modelled as a sixteen-carbon palmitic acid moiety, while the density extending from Cys378 is shorter, accommodating only a seven carbon fatty acyl chain. S-acylation occurs through the reversible covalent addition of a fatty acid to the sulfhydryl group of a cysteine residue. Both modified cysteine residues align with the cytosolic leaflet of the membrane bilayer inside the c-ring (**Fig. 2C**). S-acylation was also detected at tonoplast-specific subunit a3 biochemically, using a biotin switch assay. In this assay, acyl groups are cleaved from S-acylated proteins with hydroxylamine, replaced by biotin, and captured with neutravidin beads (Hemsley et al., 2008). Immunoblotting with an anti-GFP antibody is subsequently used to detect the protein of interest, which is only captured by the neutravidin beads if it was S-acylated. S-acylated proteins can be detected when hydroxylamine is included in the protocol (hydroxylamine +) but not when it is omitted (hydroxylamine -). A loading control, obtained by taking a sample of protein before affinity purification, is used to ensure equal loading of the samples. S-acylation of the tonoplast-specific a3 subunit (VHA-a3-GFP) was detected in hydroxylamine-treated samples (∼130 kDa, hydroxylamine +), but not the untreated controls (hydroxylamine -) (**Fig. 2D**, *upper*). No signal was detected for the TGN/EE-specific a1 subunit (VHA-a1-GFP), demonstrating that S-acylated cysteines are not conserved between the isoforms (**Fig. 2D**, *upper*). Three cysteines are present in subunit a3 but not subunit a1: Cys10, Cys11, and Cys 653 (**Fig. 2D**, *lower*). Density corresponding to S-acylation is not apparent for Cys653 in the cryo-EM map (**Fig. S3B**). Density for Cys10 and Cys11, located in the flexible N-terminal linker of subunit a3, is not visible in the cryo-EM map, preventing direct visualization of the modifications. Therefore, we mutated both Cys10 and Cys11 to serine in subunit a3 (independent lines VHA-a3SS-GFP #9 and #10) and repeated the biotin switch assay for the mutants. No signal was detected for either hydroxylamine-treated (hydroxylamine +) or untreated (hydroxylamine -) samples (**Fig. 2D**), revealing that one or both of Cys10 and Cys11 are S-acylated in the tonoplast-specific a3 subunit.

### Endogenous OXR5 interacts with V-ATPase in the TGN/EE

Approximately 3% of the protein complexes identified in cryo-EM images contributed to a 3D class that showed a density in addition to the known V-ATPase subunits (**Fig. 3A**, **Fig. S2**, and **Table S1**). This low number of particle images resulted in a cryo-EM map with substantial background noise and limited resolution, and consequently the identity of the interacting protein could not be determined directly from the map **(Fig. S1F)**. However, mass spectrometry indicated that the purified Arabidopsis V-ATPase sample contains OXidation Resistance 5 protein (OXR5). An AlphaFold3 model (Jumper et al., 2021) of the OXR5 structure (AF-Q9FKA3-F1) fit the density with high fidelity (**Fig. 3B**), identifying the additional density as OXR5. OXR5 is a TLDc-domain-containing protein (Colombatti et al., 2019; Volkert et al., 2000).

**Figure 3.**
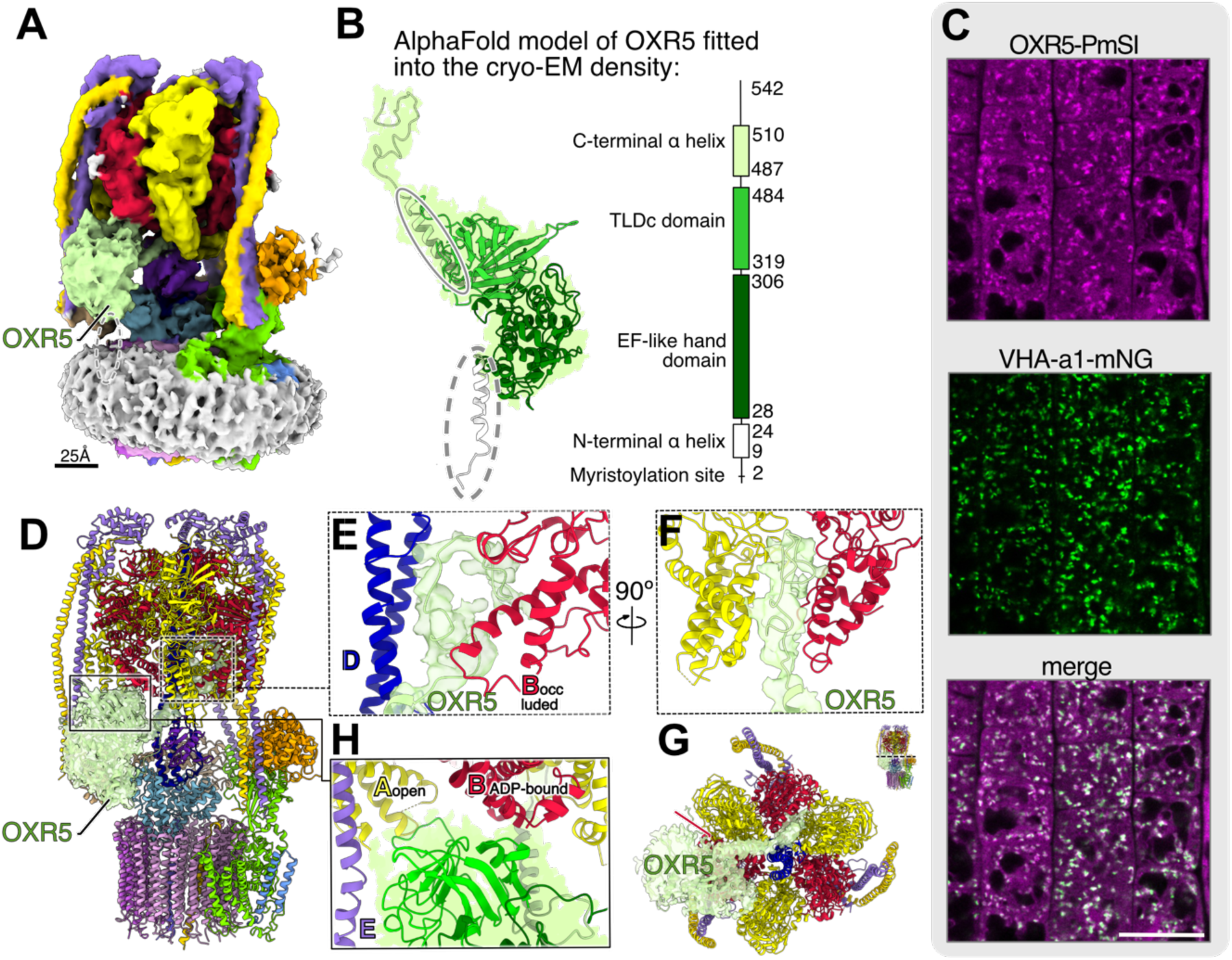
V-ATPase from the TGN/EE interacts with the TLDc protein OXR5. (A) Composite map of Arabidopsis V-ATPase in rotational state 1 bound to OXR5 (*green density*). (B) AlphaFold3 model (*left*) and domain organisation (*right*) of OXR5. (C) VHA-a1-mNG is localized at the TGN/EE. OXR5-PmScarlet-I shows partial overlap with VHA-a1-mNG at the TGN/EE. Scale bar, 20 µm. Root tip elongation zone of 5-day-old Arabidopsis seedlings were analyzed by confocal laser scanning microscopy. (D-H) Atomic model of Arabidopsis V-ATPase interactions with OXR5.

To determine the intracellular localization of OXR5, we performed confocal laser scanning microscopy in Arabidopsis lines co-expressing OXR5 fused to PmScarlet-I (OXR5-PmSI) (**Fig. 3C**, *top*) together with subunit a1, a component of V-ATPase complexes that localize to the TGN/EE, tagged with mNeonGreen (VHA-a1-mNG; **Fig. 3C**, *middle*). This experiment revealed partial co-localization of OXR5-PmSI with VHA-a1-mNG at the TGN/EE (**Fig. 3C**, *bottom*). Overexpressed OXR5-PmSI also displayed a broader endomembrane distribution as well as a cytosolic pool (**Fig. 3C**, *top*). However, OXR5-PmSI was not found at the tonoplast, indicating that it is primarily bound to V-ATPase in the TGN/EE. This restricted co-localization of OXR5-PmSI with TGN/EE-resident V-ATPases agrees with cryo-EM data showing that only approximately 3% of all V-ATPase particles are associated with OXR5. The TGN/EE V-ATPase complexes to which OXR5 binds are a small subset of the cell’s total pool of V-ATPase, which resides predominantly in the tonoplast (Carter et al., 2004; Jaquinod et al., 2007).

### OXR5 binds Arabidopsis V-ATPase similarly to mEAK-7 and Rtc5p binding of other V-ATPases

A polypeptide backbone model was constructed for OXR5 bound to V-ATPase using the amino acid sequence of the TGN-specific isoforms for the proton pump subunits **(Fig. 3D**, **Table S2)**. Like mammalian mEAK-7 and yeast Rtc5p, OXR5 contains two folded domains: an EF-hand-like domain followed by a TLDc domain, flanked by N- and C-terminal α helices (**Fig. 3B**, *ovals*). The protein’s C-terminal α helix reaches across V_1_ to interact with subunit D from the central rotor **(Fig. 3E)** and the occluded catalytic AB pair **(Fig. 3F** and **G)**. The TLDc domain of the protein binds the adjacent subunit B (ADP-bound), subunit E from the peripheral stalk, and subunit A of the open AB pair **(Fig. 3H).** The location of OXR5 binding, and the way it interacts with the V-ATPase, is similar to the interactions reported previously between V-ATPase and the OXR5 homologues mEAK-7 and Rtc5p in mammals and yeast, respectively (Khan et al., 2025; Tan et al., 2022a; L. Wang et al., 2022; R. Wang et al., 2022). The interaction is different from the binding modes of the yeast TLDc protein Oxr1p with V_1_ (Khan et al., 2022) and mammalian TLDc protein NCOA7b with V_1_ and intact V-ATPase (PDB 7UZJ, 7UZK, 7UZG, and 7UZI). Interestingly, in different organisms, OXR5 homologues interact with V-ATPase in different rotational states: the plant OXR5 and yeast Rtc5p bind rotational state 1, while mammalian mEAK-7 binds rotational state 2. However, the C-terminal α helices of OXR5, Rtc5p, and mEAK-7 all interact with the occluded catalytic AB pair from the V_1_ region. In the cryo-EM map, the N-terminal region of OXR5 is poorly resolved (**Fig. 3B**, *dashed oval*). The N-terminal α helix of OXR5 contains a conserved myristoylation motif (Gly2-Ala3-Ser4-Ser5-Ser6-Asp7-Asp8), where the Gly residue is covalently modified with myristic acid (Yamauchi et al., 2010). Myristoyl groups are typically embedded in the lipid bilayer and here would anchor OXR5 in the TGN/EE membrane.

### OXR5 binding modifies V-ATPase conformation

Binding of OXR5 to the Arabidopsis V-ATPase leads to conformational changes both in the V_1_ and V_O_ regions of the complex (**Fig. 4A**). Most strikingly, the C terminus of OXR5 binds between the central rotor subunit D and subunit B from the occluded AB pair, forcing them apart (**Fig. 4B**). This interaction also shifts the B subunit away from its corresponding A subunit, widening the cleft between the proteins (**Fig. 4C** *blue arrow*). The occluded catalytic AB pair does not interact with ATP or ADP. However, OXR5 binding may interfere with the rotary catalytic cycle of the proton pump. While OXR5 does not interact directly with the V_O_ region of the Arabidopsis V-ATPase, its binding changes the position of the rotor in the subcomplex (**Fig. 4D**). The interaction of the C-terminal α helix of OXR5 and subunit F **(Fig. 4D**, *dashed rectangle*) leads to rotation of the rotor subcomplex in the opposite direction from the rotation that occurs during proton pumping (**Fig. 4E**, *red and green arrows*). Overall, binding of OXR5 to rotational state 1 of V-ATPase changes the conformation of the state to make it more closely resemble rotational state 3. These conformational changes appear incompatible with the enzyme’s normal activity. However, mEAK-7 (Tan et al., 2022a) and Rtc5p (Khan et al., 2025) were not found to inhibit V-ATPase activity. The substoichiometric binding of OXR5 to the Arabidopsis V-ATPase means that, even if OXR5 binding was inhibitory, its binding would be unlikely to have a large effect on overall V-ATPase activity in the cell.

**Figure 4.**
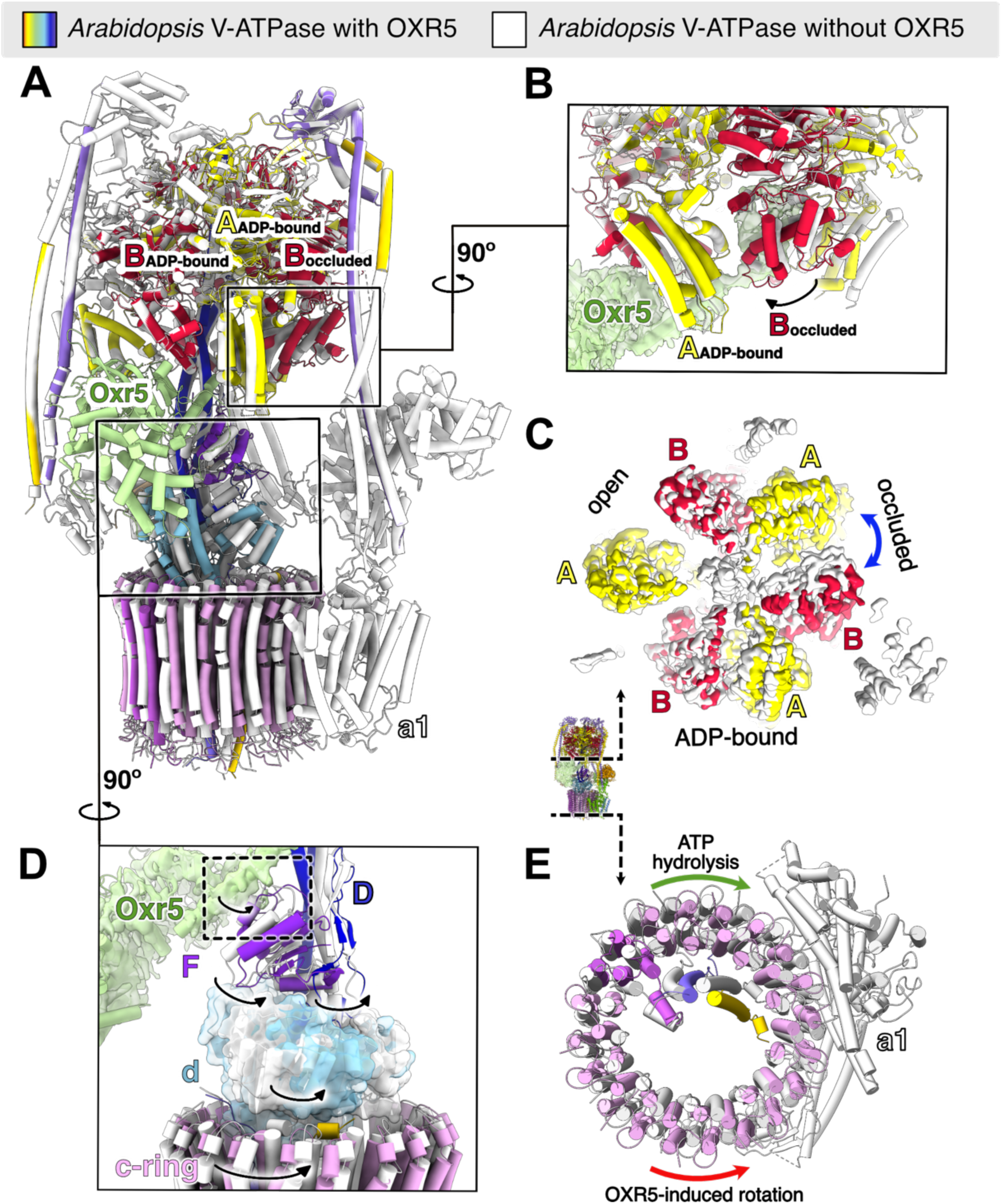
OXR5 binding causes conformational changes in *Arabidopsis* V-ATPase. (A) Overlayed models of unbound (*white*) and OXR5-bound (*coloured*) V-ATPases. (B and C) The effect of OXR5 binding on V-ATPase subunits. The interaction distorts the occluded AB pair (D) and alters the rotation of the c-ring (E).

## Discussion

Differences between the V-ATPase complexes from different organisms hint at variability in how these proton pumps are regulated. The cryo-EM structure of Arabidopsis V-ATPase presented here reveals unique post-translational modifications as well as a newly identified V-ATPase-interacting protein in plants. Further, subunit H, which is critical for inhibiting ATP hydrolysis following dissociation of V_1_ from V_O_, showed differences in conformation from the citrus V-ATPase (Tan et al., 2022b). The vacuoles of citrus fruit cells are hyperacidified (pH ∼2) owing to the activity of P-type ATPases found in the membranes of those organelles but not found in the Arabidopsis tonoplast (Strazzer et al., 2019). The novel conformation of the citrus V-ATPase was proposed to be important for preventing reverse proton transport from the lumen of the vacuole to the cytosol through the V-ATPase (Tan et al., 2022b). Consistent with this hypothesis, the Arabidopsis V-ATPase, which does not have an unusually acidic vacuole (pH 5.8), has the same conformation of the H subunit seen in yeast and mammalian V-ATPases rather than the conformations found in the citrus V-ATPase.

The S-acylation detected in the AP1 and a3 subunits occurs in ∼6% of the Arabidopsis proteome (Kumar et al., 2022). Catalyzed by one of 24 different S-acyl transferases (Batistic, 2012), S-acylation can anchor proteins to membranes, stabilize membrane protein structures, or affect intermolecular interactions to regulate protein localization and activity (Batistic et al., 2008; Sorek et al., 2010; Konrad et al., 2014; Chen et al., 2021). In the assembled Arabidopsis V-ATPase or V_O_ complex, the S-acylated cysteine residues of AP1 would be inaccessible to other enzymes, implying that the protein is modified before the c-ring is assembled around it. In contrast, the S-acylated subunit a3 is expected to be able to interact with lipids and other proteins. Therefore, the functions of S-acylation of AP1 and a3 may be different from each other. For AP1, the two acylated cysteine residues in the protein align with the cytosolic leaflet of the lipid bilayer inside the c-ring. This acylation may help position the phospholipids inside the c-ring, making the pore of the ring impermeable to protons and other ions. The experiments presented here show the acylation of tonoplast-specific subunit a3, but not the TGN/EE-specific subunit a1. The two S-acylated cysteine residues in subunit a3 are also found in the tonoplast-specific Arabidopsis a2 subunit and are conserved in both proteins in different plant species (**Fig. S4**, *red*). S-acylation of mammalian V-ATPase subunit a1 is important for targeting V-ATPase to the lysosome (Bagh et al., 2017). While the predicted modification site differs between mammalian and plant a subunits, the role of S-acylation in V-ATPase localization may be conserved.

The structure of the Arabidopsis V-ATPase bound to OXR5 provides the first evidence for interaction of TLDc proteins with V-ATPase in plants. The Arabidopsis TLDc protein OXR2 has been associated with resistance to oxidative stress (Colombatti et al., 2019) and infection by bacterial pathogens (Mencia et al., 2020) through transcriptional upregulation in these conditions. OXR5 may provide resistance to abiotic or biotic stressors through V-ATPase. Besides its TLDc domain, OXR5 includes an EF-hand-like domain and a myristoylation motif, similar to mammalian mEAK-7 (Tan et al., 2022a; L. Wang et al., 2022; R. Wang et al., 2022) and yeast Rtc5p (Khan et al., 2025) (**Fig. S5A**). The structure of OXR5 shows a cluster of cysteine residues in the core of its EF-hand-like domain (**Fig. S5B**) that could relate to potential oxidation of the protein in some conditions. OXR5 colocalizes with V-ATPases at the TGN/EE, unlike its mammalian and yeast homologues that are found at the lysosomal and vacuolar membranes, respectively (Khan et al., 2025; Nguyen et al., 2018; Tan et al., 2022a). The mechanism by which this organelle specificity is achieved for OXR5 is not clear because in the structure OXR5 does not appear to interact directly with the TGN-specific a1 subunit of V-ATPase. While binding of OXR5 to Arabidopsis V-ATPase causes conformational changes in the latter, the biological function of the interaction is not apparent from the structure. Functional studies of the mammalian and yeast homologues of OXR5 did not assign a definitive function to either of those proteins. *In vitro*, binding of mEAK-7 to V-ATPase showed no significant effect (Tan et al., 2022a) or a slight enhancement (Oot and Wilkens, 2024) of the ATPase activity of the enzyme. Recent experiments suggest that, *in vitro*, yeast Rtc5p can facilitate V-ATPase assembly (Khan et al., 2025). However, over-expression of mEAK-7 does not affect lysosomal pH (Tan et al., 2022a) and neither over-expression nor deletion of Rtc5p affects vacuolar pH (Klössel et al., 2024). Therefore, the role of OXR5 and its homologues remains unclear. Overall, the Arabidopsis V-ATPase structure presented here gives insight into the structure and regulation of these proton pumps in plants.

## Materials and Methods

### SidK expression and purification

SidK was expressed and purified by the EMBL protein expression core facility, Heidelberg, from the plasmid SidK1-178-3×FLAG (Addgene #175787) as described previously (Abbas et al., 2020). The final SidK-3×FLAG protein was prepared in a buffer with 50 mM tris-HCl, pH 7.5, 300 mM NaCl, and 10% (v/v) glycerol.

### Purification of V-ATPases from Arabidopsis thaliana Col-0

All steps were performed at 4 °C. Microsomal membranes were isolated from Arabidopsis rosette leaves (5-6 weeks old, short-day conditions: 8 h light/16 h dark). Approximately 100 g of leaf material was homogenized in ice-cold homogenization buffer (350 mM sucrose, 70 mM tris-HCl pH 7.5, 10 % (v/v) glycerol, 3 mM ethylenediaminetetraacetic acid (EDTA), 0.15 % (w/v) bovine serum albumin, 1.5 % (w/v) polyvinylpyrrolidone-40, 4 mM dithiotreitol, 1× Roche complete protease inhibitor cocktail) using a blender and subsequently filtered through two layers of Miracloth. The filtrate was clarified by centrifugation at 10,000 × g for 25 min, and the resulting supernatant was subjected to ultracentrifugation at 100,000 × g for 60 min. Microsomal membrane pellets were frozen in liquid nitrogen and stored at -80 °C. Frozen pellets were resuspended in solubilization buffer (50 mM 4-(2-hydroxyethyl)-1-piperazineethanesulfonic acid (HEPES) pH 7.0, 320 mM sucrose, 300 mM NaCl, 10% (v/v) glycerol, 5 mM EDTA, 5 mM aminocaproic acid, 5 mM para aminobenzamidine, 0.2 mM phenylmethylsulfonyl fluoride) without detergent (10 mL per g pellet), and homogenized thoroughly using a Dounce homogenizer. Membranes were solubilized by incubation with 1% (w/v) n-dodecyl-β-D-maltoside (DDM) for 2 h at 4 °C with gentle stirring. Insoluble material was removed by centrifugation at 133,500 × g for 60 min. The supernatant was retained for affinity purification.

Anti-FLAG affinity resin (Sigma-Aldrich A2220) was equilibrated with protein wash buffer 1 (50 mM tris-HCl pH 7.4, 150 mM NaCl, 0.01% (w/v) DDM). SidK-3×FLAG was loaded onto the column, and the flow-through was reapplied three times to maximize binding. For each 1 ml of anti-FLAG affinity resin, 0.6 mg SidK was applied. Solubilized microsomal membrane extract was then applied to the SidK-loaded resin, and the flow-through was reapplied to maximize binding. After washing with 10 column volumes of wash buffer 2 (50 mM HEPES pH 7, 300 mM NaCl, 0.025 % (w/v) glycodiosgenin), bound V-ATPase complexes were eluted in three column volumes of wash buffer supplemented with 150 µg/mL 3×FLAG peptide, incubated for 30-60 min before elution. Afterwards another 5 column volumes of wash buffer 2 were applied to elute remaining complexes. Purity of the eluate was evaluated with SDS-PAGE and the eluate was concentrated with a 100 kDa MWCO centrifugal concentrator and loaded on a Superose 6 Increase 10/300 gel filtration column (GE healthcare) equilibrated with wash buffer 2. The fraction corresponding to fully assembled V-ATPase was frozen in liquid nitrogen.

### Cryo-EM specimen preparation, imaging, and image analysis

Homemade holey gold grids (Marr et al., 2014) were glow discharged for 2 min in air prior to sample application. Purified protein (2 μL) at a concentration of ∼2 mg/mL (as estimated by absorbance at 280 nm) was applied onto a grid held in the tweezers of an EM GP2 grid freezing device (Leica) at 4 °C and ∼90% relative humidity. Grids were blotted for 2 s before plunge freezing in liquid ethane. The grids were screened with a Glacios 2 electron microscope operating at 200 kV and equipped with a Falcon 4i camera (Thermo Fisher Scientific). High resolution data collection was then performed with a Titan Krios G3 electron microscope operating at 300 kV and equipped with a Falcon 4i camera and a Selectris X energy filter (Thermo Fisher Scientific) using a slit width of 10 eV. Electron microscopy data acquisition was automated with the EPU software (Thermo Fisher Scientific). For high resolution data collection, 7681 movies were collected in electron event representation mode (Guo et al., 2020) at 130,000× nominal magnification, corresponding to a calibrated pixel size of 0.93 Å, with a total exposure of ∼40 e^−^/Å^2^. Data quality was monitored in real time with cryoSPARC Live (Punjani et al., 2017).

### Cryo-EM image analysis

Cryo-EM data analysis was performed in cryoSPARC (Punjani et al., 2017). Exposures were aligned, and contrast transfer function (CTF) parameters estimated in patches. Initially, particle images from a subset of 175 micrographs were manually selected and extracted with a box size of 512 × 512 pixels (0.93 Å/pixel) and subjected to two-dimensional (2D) classification. The best 2D class averages were then used as templates for particle selection on the same subset of micrographs. Particle images found with templates were cleaned by several rounds of 2D classification and then were used to train a Topaz model (Bepler et al., 2019). The model was applied to the whole dataset and selected 190,490 particle images, which were extracted with a box size of 512 × 512 pixels. The selected particle images were aligned with local motion correction (Rubinstein and Brubaker, 2015) followed by several rounds of cleaning with 2D classification. The selected particle images were then cleaned and divided into three rotational states with multiple iterations of ab initio reconstruction and heterogeneous refinement. Three high-resolution maps were calculated with homogeneous refinement from 144,507 particle images. Individual particle CTF parameters were estimated for the pooled set of particle images. The corrected particles were once again divided into the three 3D maps by ab initio reconstruction and heterogeneous refinement, resulting in three sets of particle images: 54,020 particle images for rotational state 1; 48,315 particle images for rotational state 2; and 42,172 particle images for rotational state 3. Each dataset underwent an additional round of homogeneous refinement prior to local refinements of the V_1_ and V_O_ regions in each rotational state. 3D classification with focused masks was used to further clean maps for each subcomplex. For the regions of the map that do not move between rotational states (subunits a, C, and the bottom halves of the peripheral stalks), local refinement was performed on the pooled set of particle images aligned to the 3D map of rotational state 1.

Particle images with OXR5 bound to V-ATPase were initially separated by 3D classification without a focused mask from the dataset corresponding to rotational state 1. 3D classification with the focused mask around the OXR5 density was then used to clean the subset of images to 5,323 particle images. A map of V-ATPase bound to OXR5 was calculated with ab initio reconstruction. Homogeneous refinement was performed to obtain the overall map of V-ATPase and local refinements were performed to obtain the maps of the V_1_ and V_O_ regions. The overall map had a nominal resolution of 4.3 Å, with local resolution reaching 3.6 Å for the A_3_B_3_DFd region and 4.0 Å for the c_9_c″a region. However, many regions of the map appear to be at substantially lower resolution.

### Model building

An initial atomic model for rotational state 1 of V-ATPase was generated with ModelAngelo (Jamali et al., 2024) using the amino acid sequences of the V-ATPase subunits as determined by mass spectrometry. Large gaps in the model of subunits a, D, F, and C were filled with segments of proteins obtained from the AlphaFoldDB protein structure database (Abramson et al., 2024) and fit into the maps as rigid bodies with UCSF ChimeraX (Goddard et al., 2018). Models for rotational states 2 and 3 of V-ATPase, as well as the model of the OXR5 bound complex, were created from the model of rotational state 1 by rearrangement of subunits in UCSF ChimeraX. Models for OXR5 and the a1 subunit were obtained from the AlphaFoldDB database (AF-Q9FKA3-F1 and AF-Q8RWZ7-F1, respectively). Combining model fragments, optimization of model-to-map fit, and molecular dihedral angles, filling of small gaps in the starting model, and fitting of ADP and lipid modifications were done with Coot v0.9.6 (Emsley et al., 2010) and ISOLDE v1.8 (Croll, 2018). Models were further refined with PHENIX v1.19.2 (Adams et al., 2010). All images of molecular structures were rendered with UCSF ChimeraX v1.8.

### Mass spectrometry

Microsomal proteins were predigested for three hours with endoproteinase Lys-C (0.5 µg/µL; Wako Chemicals, Neuss) at room temperature. After 4-fold dilution with 10 mM tris-HCl (pH 8), samples were digested with 4 µL sequencing grade modified trypsin (0.5 µg/µL; Promega) overnight at 37 °C. Following overnight digestion, trifluoroacetic acid (TFA) was added (until pH ≤ 3) to stop digestion. Digested peptides were desalted with C18 tips (Rappsilber et al., 2003) and dissolved in 100 µL 80% (v/v) acetonitrile and 0.1% (v/v) trifluoroacetic acid before LC-MS/MS analysis. Tryptic peptide mixtures were analyzed by LC/MS/MS using nanoflow Easy-nLC1000 (Thermo Scientific) as an HPLC-system and a Quadrupole-Orbitrap hybrid mass spectrometer (Q-Exactive Plus, Thermo Scientific) as a mass analyzer. Peptides were eluted from a 75 μm × 25 cm C18 analytical column (PepMan, Thermo Scientific) on a linear gradient running from 4 to 64% acetonitrile in 70 min and sprayed directly into the Q-Exactive mass spectrometer. Proteins were identified by MS/MS using information-dependent acquisition of fragmentation spectra of multiple charged peptides. Up to twelve data-dependent MS/MS spectra were acquired for each full-scan spectrum acquired at 70,000 full-width half-maximum resolution with an ion target of 1×10^6^. Fragment spectra were acquired at a resolution of 35,000 with an ion target 1×10^5^. Overall cycle time was approximately 1 s.

Protein identification and ion intensity quantitation was carried out by MaxQuant version 2.2.0.0 (Cox and Mann, 2008). Spectra were matched against the *A. thaliana* proteome (TAIR10, 35386 entries) using Andromeda (Cox et al., 2011). Carbamidomethylation of cysteine was set as a fixed modification; oxidation of methionine as well as phosphorylation of serine, threonine and tyrosine was set as variable modifications. Mass tolerance for the database search was set to 20 ppm on full scans and 0.5 Da for fragment ions. Multiplicity was set to 1. For label-free quantitation, retention time matching between runs was chosen within a time window of two minutes. Peptide false discovery rate (FDR) and protein FDR were set to 0.01, while site FDR was set to 0.05. Hits to contaminants (e.g. keratins) and reverse hits identified by MaxQuant were excluded from further analysis. The option of calculating iBAQ values (Schwanhäusser et al., 2011) was selected to later allow for comparison of protein abundances within each sample. Protein data was analyzed based on the proteingroups.txt output from MaxQuant according to their iBAQ values.

### Construct design for confocal microscopy experiments and biotin switch assay

For preparation of UBQ:OXR5-PmSI plasmid the GreenGate cloning system was employed (Lampropoulos et al., 2013). The full-length coding sequence of OXR5 (AT5G39590) lacking the stop codon was amplified from Col-0 cDNA using primers 5383-fw and 5384-rv (**Supplementary Data 2**). Following Eco31I digestion, the 1626 bp fragment was cloned into pGGC000. The complete UBQ:OXR5-PmSI construct was assembled from the GreenGate modules pGGA UBQ10 promoter, pGGB B-dummy, pGGC OXR5, pGGD GSL-PmSI, pGGE tHSP18.2M, and pGGF pMAS:SulfRm:t35s in pGGZ004 (Lampropoulos et al., 2013; Lupanga et al., 2020). The final construct was transformed in Arabidopsis lines expressing VHA-a1-mNeonGreen (VHA-a1-mNG) (McKay et al., 2025). For generation of AP1-mNG, the full-length genomic sequence of AP1 (At3g13410) lacking the stop codon was amplified from Col-0 gDNA using primers 3092-fw and 3095-rv (**Supplementary Data 2**). Following Eco31I digestion, the 1919 bp fragment was cloned into pGGC000. The complete VHA-A-AP1-mNG construct was assembled from the GreenGate modules pGGA UBQ10 promoter, pGGB B-dummy, pGGC AP1, pGGD GSL-mNG, pGGE tHSP18.2M, and pGGF pMAS:SulfRm:t35s in pGGZ004 (Lampropoulos et al., 2013). Constructs were transformed into Col-0 and into the *vha-a2 vha-a3* double mutant (Krebs et al., 2010). To obtain a fusion protein of AP2 with GFP, the At3g24160 CDS was amplified from cDNA of Col-0 seedlings with 1895 and 1937 primers, inserting the SacII restriction site N-terminally and the MluI restriction site at the C terminus. The stop codon was removed with this primer. The PCR product was cloned in pUGT1kan vector. The construct was transformed into Col-0 and into the *vha-a2 vha-a3* double mutant (Krebs et al., 2010). For the generation of VHA-a3SS-GFP the two conserved cysteine residues at the N terminus of VHA-a3 were substituted with serines using a two-step PCR strategy with overlapping primers VHA-a3SS-GFP PCR1 forward and reverse. In the first PCR, residues C10 and C11 were mutated to serine using VHA-a3 cDNA (VHA-a3 in pBluescript) as the template. The resulting PCR product was then used as the template for a second PCR, which introduced an *Apa*I restriction site at the 5′ end and a *Spe*I site at the 3′ end with primers VHA-a3SS-GFP PCR2 forward and reverse. The final PCR product was blunt-end cloned into the pJET1.2 vector and verified by sequencing. A confirmed clone was subsequently digested with *Apa*I and *Spe*I to generate compatible cohesive ends. The resulting fragment was ligated into the destination vector pUGT1Kan, a derivative of pUTkan containing GFP (S65T) for C-terminal fusion to the protein of interest (Krebs et al., 2012), which had been digested with the same enzymes. Plant transformation was performed by floral dipping (Zhang et al., 2006). Preparation of VHA-a1-GFP and VHA-a3-GFP lines was described previously (Dettmer et al., 2006).

### Confocal laser scanning microscopy

*Arabidopsis thaliana* (Col-0) seedlings were cultivated on agar plates. The growth medium consisted of half-strength Murashige and Skoog (MS) basal salts (Duchefa), supplemented with 0.5% (w/v) sucrose and 0.5% (w/v) phyto agar. The pH of the medium was adjusted to 5.8 with KOH. Seeds were surface-sterilized with ethanol and stratified for 48 h at 4 °C prior to germination. Seedlings were grown under long-day conditions (16 h light/8 h dark) for 5 days. Root cells of 5-day-old seedlings were imaged with a Leica Stellaris 8 confocal laser scanning microscope equipped with a Leica HC PL APO CS2 63×/1.20 water immersion objective. Images were acquired above the division zone. Excitation wavelengths were 495 nm for mNeonGreen, and 575 nm for PmScarlet-I, using a white light laser. Image acquisition was performed with Leica LAS X software. Post-processing (Gaussian blur 1.0; global adjustment of brightness and contrast) was conducted with Fiji (Schindelin et al., 2012).

### S-Acylation Assay

S-acylated proteins were enriched using a biotin switch assay described previously (Hemsley et al., 2008) with minor modifications. Leaf tissue from 2-4 *Arabidopsis thaliana* leaves expressing VHA-a1-GFP, VHA-a3-GFP, or VHA-a3SS-GFP (lines #9 and #10) was frozen in liquid nitrogen, ground to a fine powder, and resuspended in lysis buffer (1× phosphate-buffered saline (PBS) pH 7.4, 1× protease inhibitor cocktail, 1 mM EDTA, 1% (v/v) Triton X-100, and 25 mM freshly prepared N-ethylmaleimide). Samples were incubated for 1 h at 4 °C, and insoluble material was removed by centrifugation at 500 × g for 10 min at 4 °C.

The protein concentration of the solution was determined with a bicinchoninic acid assay kit (Thermo Scientific) and adjusted to 1 mg/ml with lysis buffer. After incubation at 4 °C with gentle mixing overnight, proteins were precipitated by methanol/chloroform extraction with three volumes methanol, one volume chloroform, and four volumes water, followed by centrifugation at 10,000 × g for 30 min at 14 °C. Protein pellets were mixed with four volumes methanol for 20 min at -20 °C, centrifuged at 5,000 × g for 20 min at 4 °C, and dried in air. These pellets were next resuspended in resuspension buffer (1× PBS, pH 7.4, 8 M urea, 2% (w/v) sodium dodecyl sulfate (SDS)) by sonication and gentle agitation at room temperature. Samples were divided into two equal aliquots. One aliquot (hydroxylamine +) was treated with 800 µL freshly prepared hydroxylamine solution (1 M hydroxylamine, 1 mM EDTA, 1× protease inhibitor cocktail, and 4 mM N-[6-(biotinamido)hexyl]-3-(2-pyridyldithio)propionamide (biotin-HPDP), while the control aliquot (hydroxylamine -) was treated with the same solution without hydroxylamine (50 mM Tris-HCl, pH 7.4, 1 mM EDTA, 1 × protease inhibitor cocktail, 4 mM biotin-HPDP). Hydroxylamine specifically cleaves thioester-linked acyl groups from cysteine residues, exposing sulfhydryl groups for labelling with biotin-HPDP. Following labelling, proteins were precipitated again by methanol/chloroform extraction.

Pellets were resuspended in 100 µL resuspension buffer and diluted with 900 µL PBS containing 0.2% (v/v) Triton X-100. A sample (50-100 µL) was removed prior to affinity purification as a loading control, methanol/chloroform precipitated, and dissolved in 2 × SDS sample buffer (125 mM Tris-HCl pH 6.6, 1% (v/v) β-mercaptoethanol, 6% SDS, 0.005% (w/v) bromophenol blue, 40% (v/v) glycerol). The remaining sample was incubated with 25 µL high-capacity NeutrAvidin agarose beads for 1 h at room temperature on a rotating wheel to enrich biotinylated proteins. Beads were collected by centrifugation at 1,000 × g and washed twice with wash buffer (1× PBS pH 7.4, 500 mM NaCl, 0.1% (w/v) SDS, 1 mM EDTA, 1× protease inhibitor cocktail). Bound proteins were eluted by boiling in 2 × SDS sample buffer for 5 min at 95 °C and analysed by SDS-PAGE and immunoblotting using an anti-GFP antibody (Roth et al. 2018).

## Author contributions

KS and JLR conceived the study. NW purified the protein, generated transgenic OXR lines, and performed confocal laser scanning microscopy. FF generated transgenic VHA-AP1 and VHA-AP2 lines. UL generated transgenic VHA-a3SS-GFP. UL and FF performed S-acylation detection experiments. MK performed the cryo-EM with guidance from HW, calculated the cryo-EM maps, and constructed atomic models. WS performed the mass spectrometry analysis. MK and JLR wrote the manuscript and prepared the figures with input from the other authors, who all contributed to editing of the manuscript.

## Acknowledgments

We thank Dr. Samir Benlekbir for assistance with collecting cryo-EM data. We thank Prof. Cristina Paulino, Dr. Dirk Flemming, and Dr. Jochen Baßler for their support with negative-stain microscopy and size exclusion chromatography of the initial V-ATPase samples. MK was supported by a ResTraComp scholarship from the Hospital for Sick Children, HW was supported by Mary H. Beatty Fellowship, and JLR was supported by the Canada Research Chairs program. This work was supported by grants PJT195707 from the Canadian Institutes of Health Research (JLR) and by the Deutsche Forschungsgemeinschaft within CRC1101 (KS). Cryo-EM data were collected at the Toronto High-Resolution High-Throughput cryo-EM Facility at The Hospital for Sick Children, supported by the Canada Foundation for Innovation and the Government of Ontario.

---

**Figure S1.**
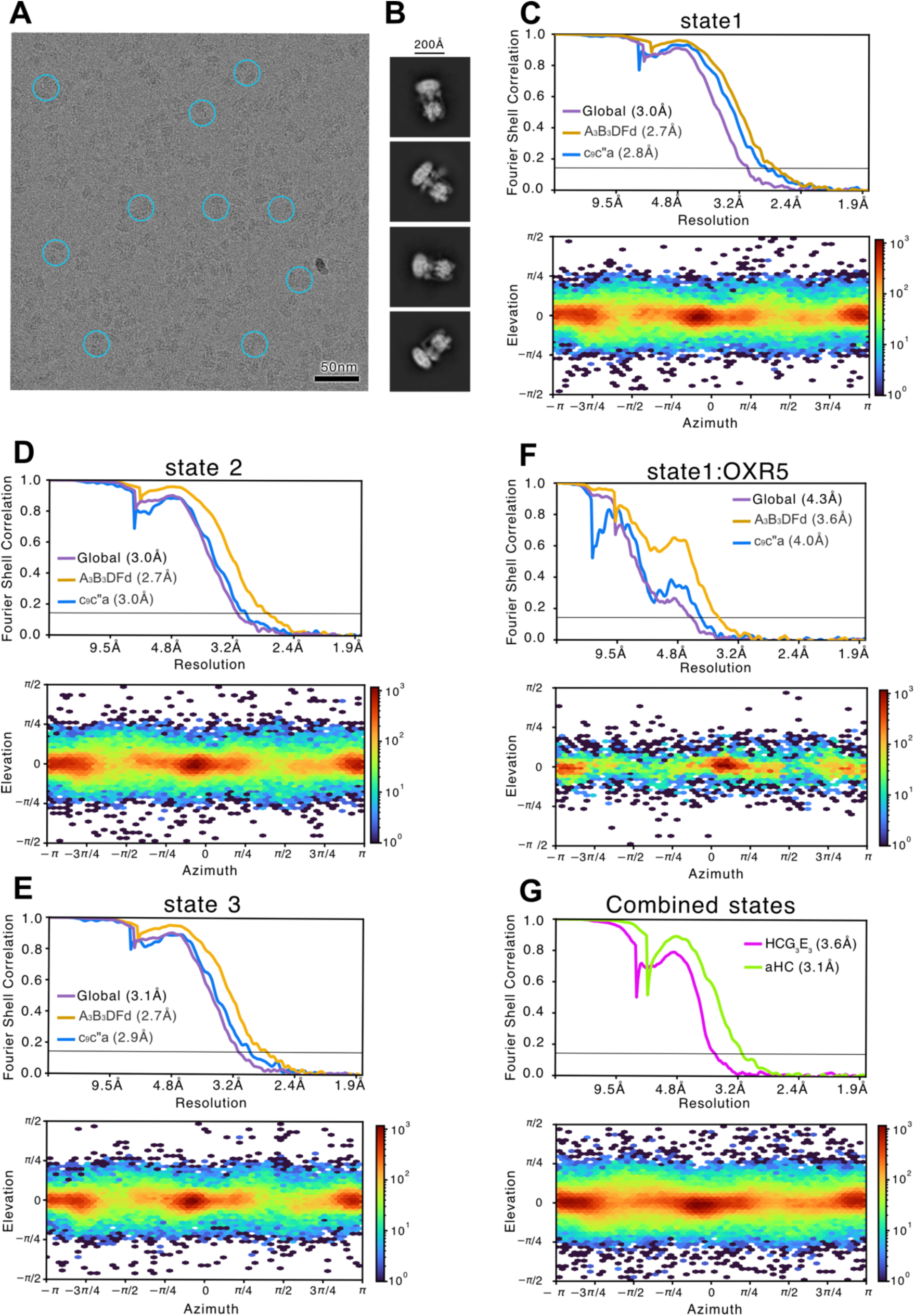
Arabidopsis V-ATPase cryo-EM validation. (A) Example micrograph. (B) Example 2D class average images. (C - G) Fourier Shell Correlation plots following a gold standard refinement and correction for the effects of masking (*top*) and orientation distribution plots (*bottom*) for state 1 (C), state 2 (D), state 3 (E), state 1 with OXR5 (F), and focused refinements from all states (G).

**Figure S2.**
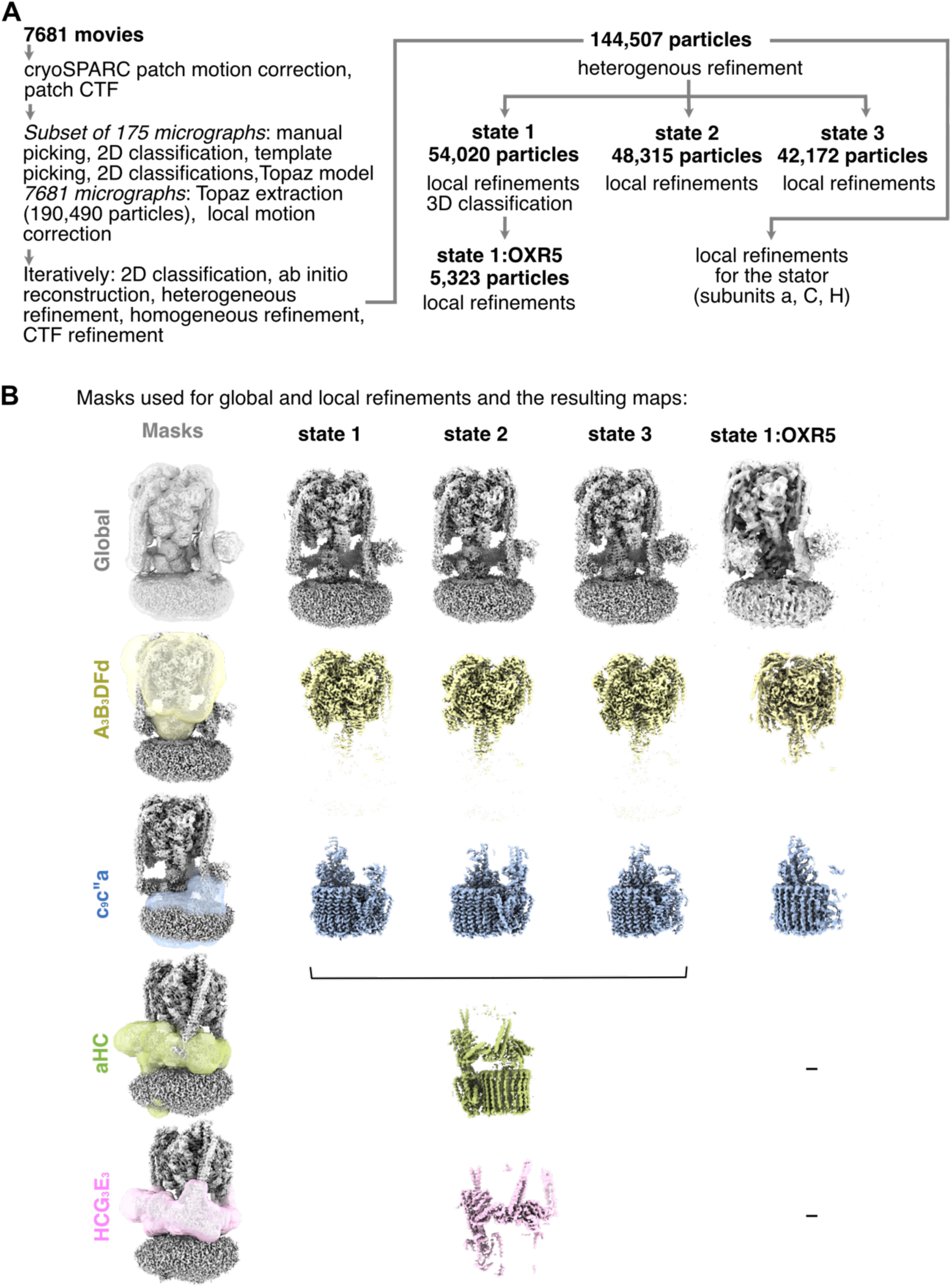
Cryo-EM workflow. (A) Workflow used to identify three rotational states and the OXR5-bound structure for Arabidopsis V-ATPase. (B) Global and local refinement schemes and the corresponding high-resolution maps.

**Figure S3.**
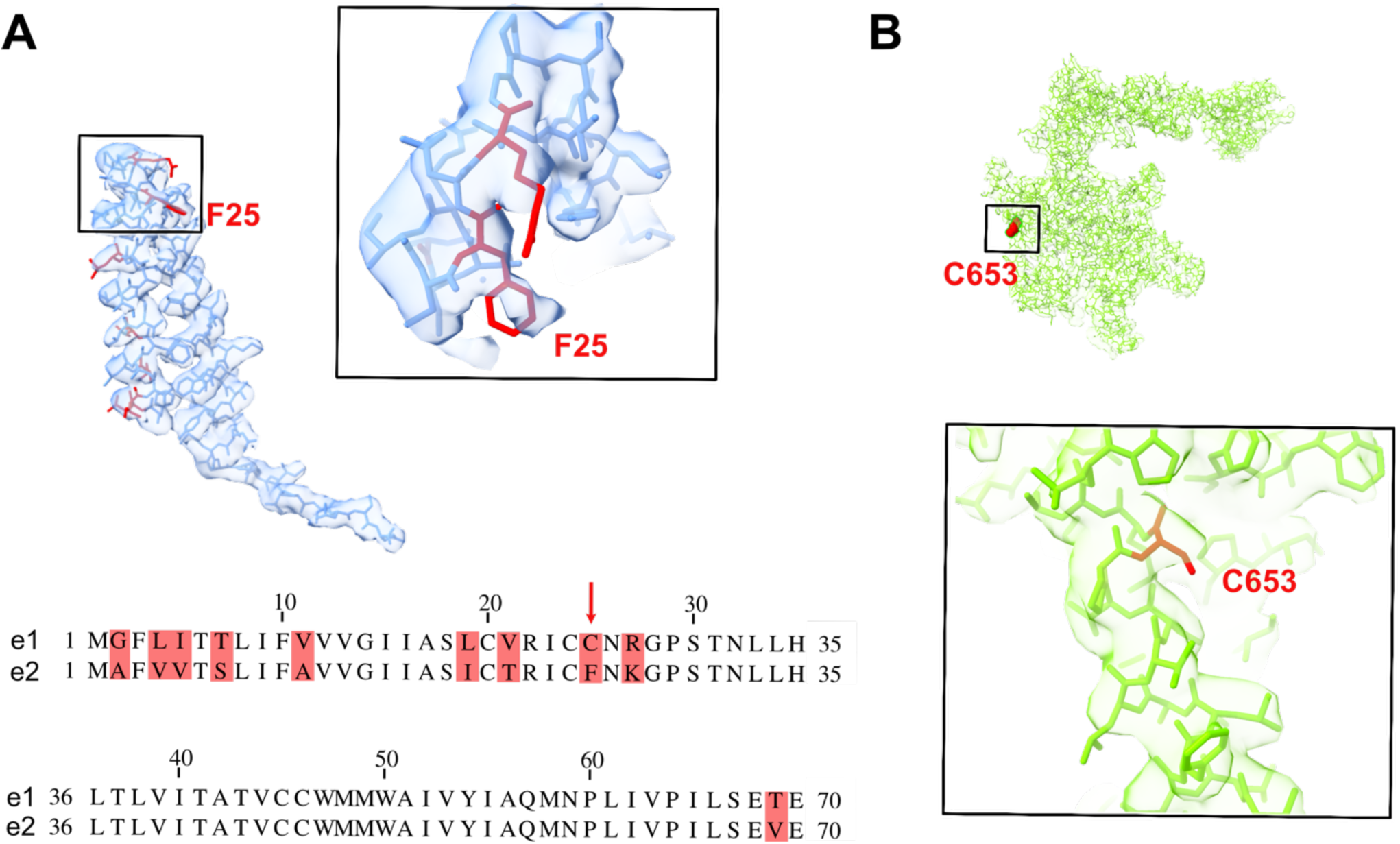
(A) Subunit e2 isoform was modeled into the cryo-EM map of Arabidopsis V-ATPase. The non-conserved residue F25 of e2 fits into the cryo-EM density. (B) No S-acylation is detected at residue C653 of subunit a3.

**Figure S4.**
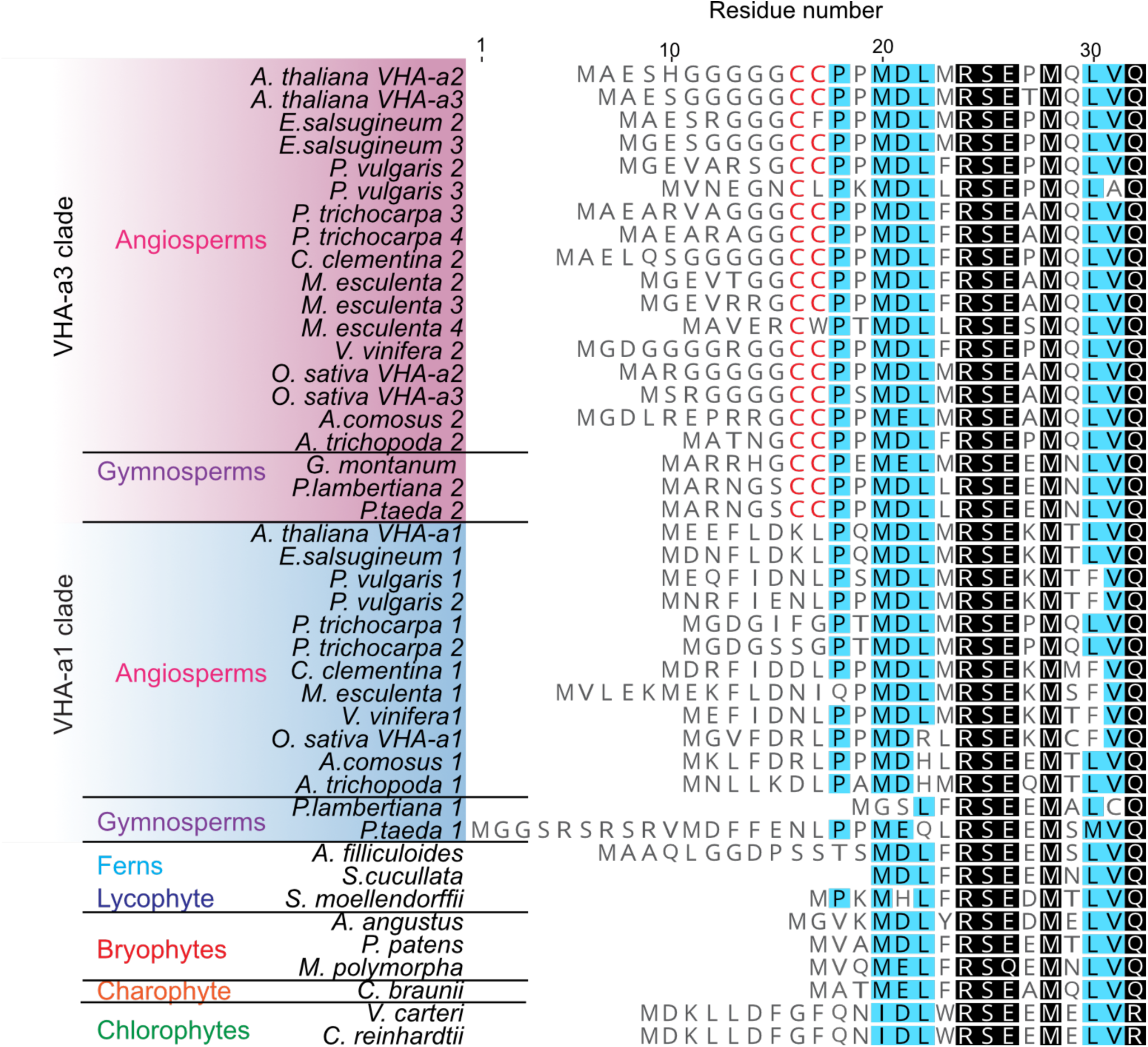
Conservation of S-acylated cysteines in different subunit a isoforms across plant species.

**Figure S5.**
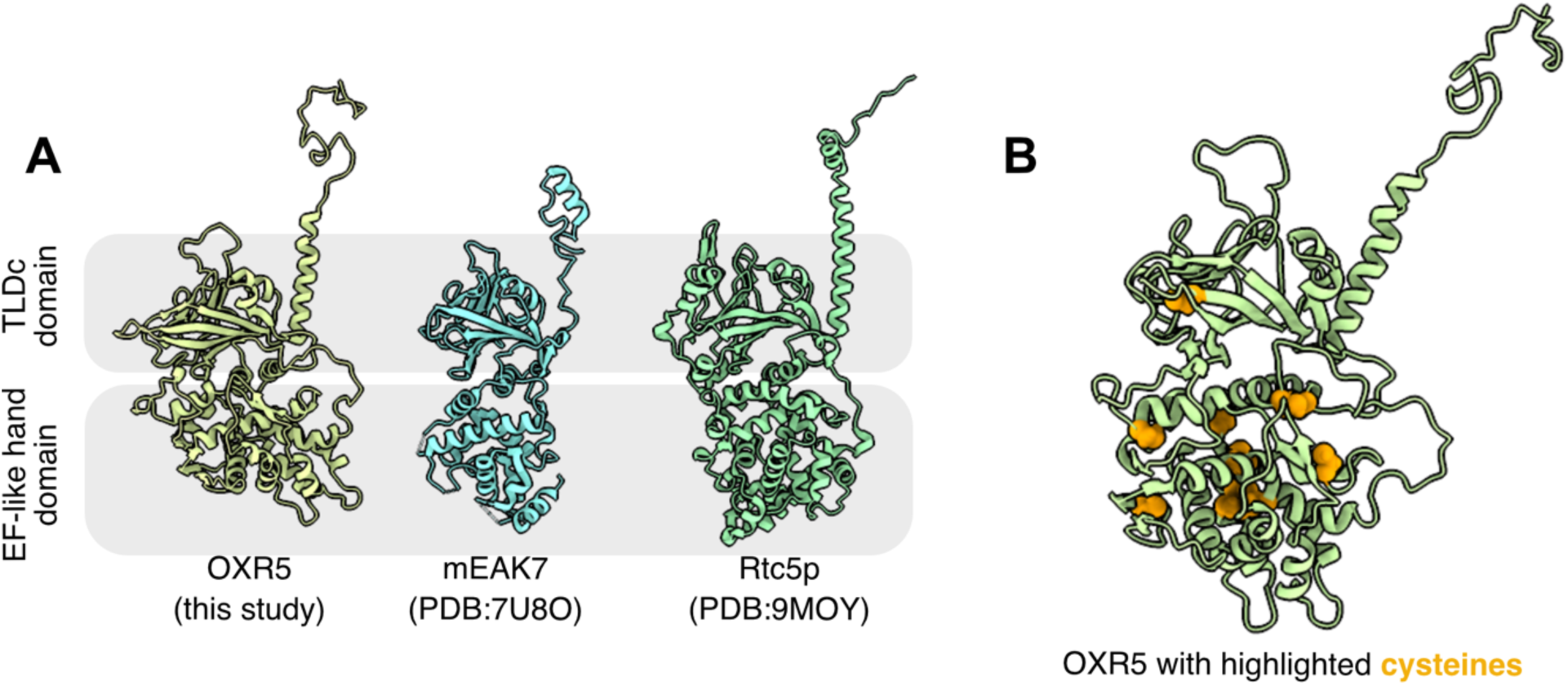
OXR5 homologues and structural features. (A) Atomic models for OXR5 and its homologues mEAK7 in mammals and Rtc5p in yeasts. (B) A cluster of twelve cysteines in OXR5. Cysteines are shown with orange spheres.

**Table S1.**
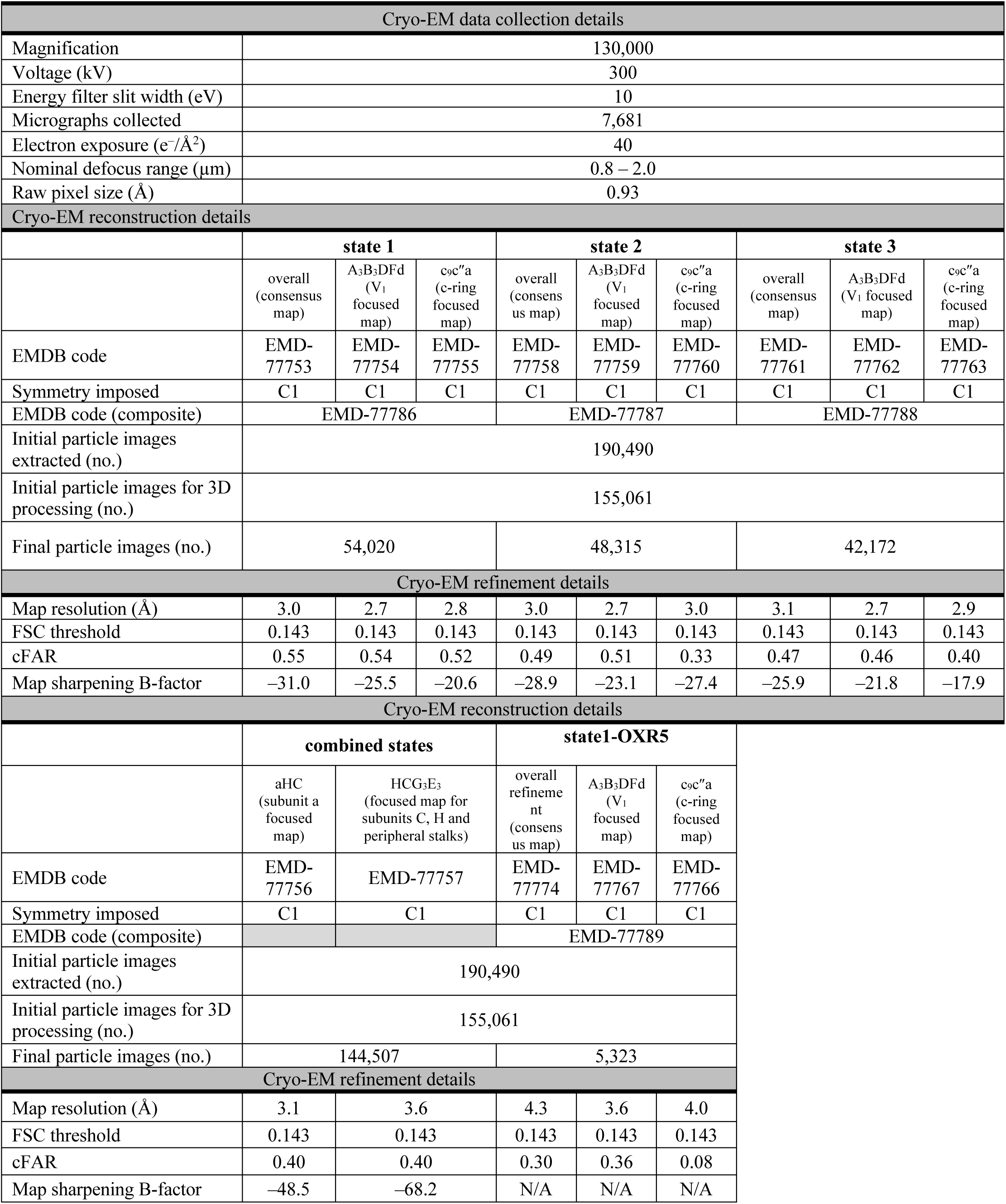
Cryo-EM map statistics.

**Table S2.**
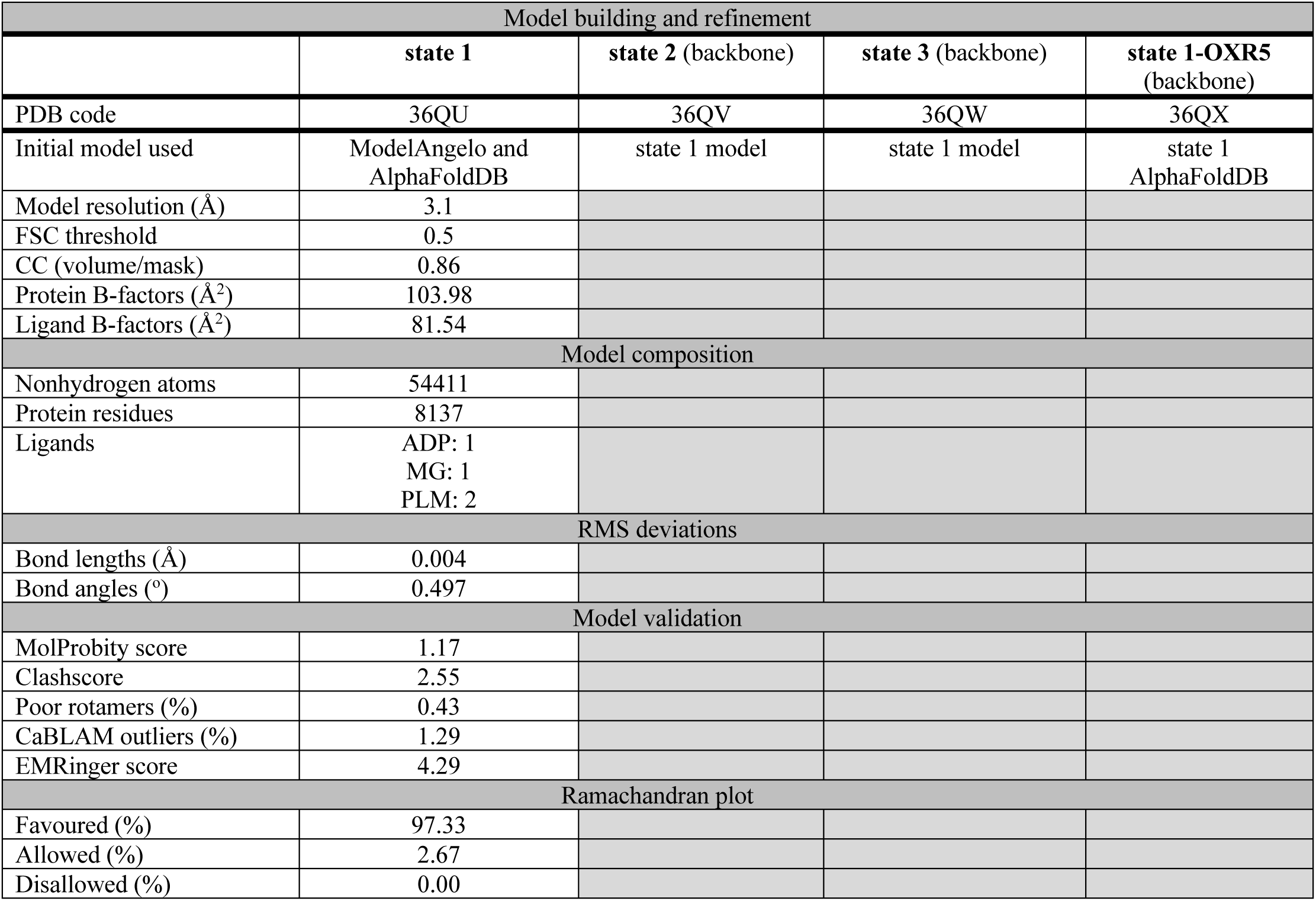
Atomic model statistics.

| Model building and refinement |  |  |  |  |
| --- | --- | --- | --- | --- |
|  | state 1 | state 2 (backbone) | state 3 (backbone) | state 1-OXR5 (backbone) |
| PDB code | 36QU | 36QV | 36QW | 36QX |
| Initial model used | ModelAngelo and AlphaFoldDB | state 1 model | state 1 model | state 1 AlphaFoldDB |
| Model resolution (Å) | 3.1 |  |  |  |
| FSC threshold | 0.5 |  |  |  |
| CC (volume/mask) | 0.86 |  |  |  |
| Protein B-factors (Å<sup>2</sup>) | 103.98 |  |  |  |
| Ligand B-factors (Å<sup>2</sup>) | 81.54 |  |  |  |
| Model composition |  |  |  |  |
| Nonhydrogen atoms | 54411 |  |  |  |
| Protein residues | 8137 |  |  |  |
| Ligands | ADP: 1<br>MG: 1<br>PLM: 2 |  |  |  |
| RMS deviations |  |  |  |  |
| Bond lengths (Å) | 0.004 |  |  |  |
| Bond angles (°) | 0.497 |  |  |  |
| Model validation |  |  |  |  |
| MolProbity score | 1.17 |  |  |  |
| Clashscore | 2.55 |  |  |  |
| Poor rotamers (%) | 0.43 |  |  |  |
| CaBLAM outliers (%) | 1.29 |  |  |  |
| EMRinger score | 4.29 |  |  |  |
| Ramachandran plot |  |  |  |  |
| Favoured (%) | 97.33 |  |  |  |
| Allowed (%) | 2.67 |  |  |  |
| Disallowed (%) | 0.00 |  |  |  |

**Supplementary Data 1. Mass spectrometry spreadsheet.**

**Supplementary Data 2. Vectors and Primers.**

## References

Abbas, Y.M., Wu, D., Bueler, S.A., Robinson, C.V., Rubinstein, J.L., 2020. Structure of V-ATPase from the mammalian brain. Science 367, 1240–1246.

Abramson, J., Adler, J., Dunger, J., Evans, R., Green, T., Pritzel, A., Ronneberger, O., Willmore, L., Ballard, A.J., Bambrick, J., Bodenstein, S.W., Evans, D.A., Hung, C.-C., O’Neill, M., Reiman, D., Tunyasuvunakool, K., Wu, Z., Žemgulytė, A., Arvaniti, E., Beattie, C., Bertolli, O., Bridgland, A., Cherepanov, A., Congreve, M., Cowen-Rivers, A.I., Cowie, A., Figurnov, M., Fuchs, F.B., Gladman, H., Jain, R., Khan, Y.A., Low, C.M.R., Perlin, K., Potapenko, A., Savy, P., Singh, S., Stecula, A., Thillaisundaram, A., Tong, C., Yakneen, S., Zhong, E.D., Zielinski, M., Žídek, A., Bapst, V., Kohli, P., Jaderberg, M., Hassabis, D., Jumper, J.M., 2024. Accurate structure prediction of biomolecular interactions with AlphaFold 3. Nature 630, 493–500. 10.1038/s41586-024-07487-w

Adams, P.D., Afonine, P.V., Bunkóczi, G., Chen, V.B., Davis, I.W., Echols, N., Headd, J.J., Hung, L.-W., Kapral, G.J., Grosse-Kunstleve, R.W., McCoy, A.J., Moriarty, N.W., Oeffner, R., Read, R.J., Richardson, D.C., Richardson, J.S., Terwilliger, T.C., Zwart, P.H., 2010. PHENIX: a comprehensive Python-based system for macromolecular structure solution. Acta Crystallogr. D Biol. Crystallogr. 66, 213–221. 10.1107/S0907444909052925

Apse, M.P., Aharon, G.S., Snedden, W.A., Blumwald, E., 1999. Salt Tolerance Conferred by Overexpression of a Vacuolar Na+/H+ Antiport in Arabidopsis. Science 285, 1256–1258. 10.1126/science.285.5431.1256

Bagh, M.B., Peng, S., Chandra, G., Zhang, Z., Singh, S.P., Pattabiraman, N., Liu, A., Mukherjee, A.B., 2017. Misrouting of v-ATPase subunit V0a1 dysregulates lysosomal acidification in a neurodegenerative lysosomal storage disease model. Nat. Commun. 8, 14612. 10.1038/ncomms14612

Batistic, O., 2012. Genomics and localization of the Arabidopsis DHHC-cysteine-rich domain S-acyltransferase protein family. Plant Physiol. 160, 1597–1612. 10.1104/pp.112.203968

Batistic, O., Sorek, N., Schültke, S., Yalovsky, S., Kudla, J., 2008. Dual fatty acyl modification determines the localization and plasma membrane targeting of CBL/CIPK Ca2+ signaling complexes in Arabidopsis. Plant Cell 20, 1346–1362. 10.1105/tpc.108.058123

Benlekbir, S., Bueler, S.A., Rubinstein, J.L., 2012. Structure of the vacuolar-type ATPase from Saccharomyces cerevisiae at 11-Å resolution. Nat. Struct. Mol. Biol. 19, 1356–62. 10.1038/nsmb.2422

Bepler, T., Morin, A., Rapp, M., Brasch, J., Shapiro, L., Noble, A.J., Berger, B., 2019. Positive-unlabeled convolutional neural networks for particle picking in cryo-electron micrographs. Nat. Methods 16, 1153–1160. 10.1038/s41592-019-0575-8

Carter, C., Pan, S., Zouhar, J., Avila, E.L., Girke, T., Raikhel, N.V., 2004. The Vegetative Vacuole Proteome of Arabidopsis thaliana Reveals Predicted and Unexpected Proteins. Plant Cell 16, 3285–3303. 10.1105/tpc.104.027078

Catalá, R., Santos, E., Alonso, J.M., Ecker, J.R., Martínez-Zapater, J.M., Salinas, J., 2003. Mutations in the Ca2+/H+ Transporter CAX1 Increase CBF/DREB1 Expression and the Cold-Acclimation Response in Arabidopsis. Plant Cell 15, 2940–2951. 10.1105/tpc.015248

Chen, D., Hao, F., Mu, H., Ahsan, N., Thelen, J.J., Stacey, G., 2021. S-acylation of P2K1 mediates extracellular ATP-induced immune signaling in Arabidopsis. Nat. Commun. 12, 2750. 10.1038/s41467-021-22854-1

Chen, J., Tang, F., Qin, L., Fang, W., Guan, L., Wu, X., Li, H., Duan, Y., Wang, F., Peng, C., Liu, Z., Wang, J., Huang, X., Wang, Lin, Yang, H., Wang, Li, Sha, W., Cai, X., Lyu, L.-D., Liu, H., Liu, F., Ge, B., Zheng, R., 2026. Mycobacterium tuberculosis modulates phosphorylation of host ATP6V1E1 to promote intracellular survival. Nat. Commun. 17, 2434. 10.1038/s41467-026-69331-1

Colombatti, F., Mencia, R., Garcia, L., Mansilla, N., Alemano, S., Andrade, A.M., Gonzalez, D.H., Welchen, E., 2019. The mitochondrial oxidation resistance protein AtOXR2 increases plant biomass and tolerance to oxidative stress. J. Exp. Bot. 70, 3177–3195. 10.1093/jxb/erz147

Coupland, C.E., Karimi, R., Bueler, S.A., Liang, Y., Courbon, G.M., Di Trani, J.M., Wong, C.J., Saghian, R., Youn, J.-Y., Wang, L.-Y., Rubinstein, J.L., 2024. High-resolution electron cryomicroscopy of V-ATPase in native synaptic vesicles. Science 385, 168–174. 10.1126/science.adp5577

Cox, J., Mann, M., 2008. MaxQuant enables high peptide identification rates, individualized p.p.b.-range mass accuracies and proteome-wide protein quantification. Nat. Biotechnol. 26, 1367–1372. 10.1038/nbt.1511

Cox, J., Neuhauser, N., Michalski, A., Scheltema, R.A., Olsen, J.V., Mann, M., 2011. Andromeda: a peptide search engine integrated into the MaxQuant environment. J. Proteome Res. 10, 1794–1805. 10.1021/pr101065j

Croll, T.I., 2018. ISOLDE: a physically realistic environment for model building into low-resolution electron-density maps. Acta Crystallogr. Sect. Struct. Biol. 74, 519–530. 10.1107/S2059798318002425

Dettmer, J., Hong-Hermesdorf, A., Stierhof, Y.-D., Schumacher, K., 2006. Vacuolar H+-ATPase Activity Is Required for Endocytic and Secretory Trafficking in Arabidopsis. Plant Cell 18, 715–730. 10.1105/tpc.105.037978

Dettmer, J., Liu, T.-Y., Schumacher, K., 2010. Functional analysis of *Arabidopsis* V-ATPase subunit VHA-E isoforms. Eur. J. Cell Biol., Mechanisms of Cell Behaviour in Eukaryotes 89, 152–156. 10.1016/j.ejcb.2009.11.008

Eaton, A.F., Brown, D., Merkulova, M., 2021. The evolutionary conserved TLDc domain defines a new class of (H+)V-ATPase interacting proteins. Sci. Rep. 11, 22654. 10.1038/s41598-021-01809-y

Emsley, P., Lohkamp, B., Scott, W.G., Cowtan, K., 2010. Features and development of *Coot*. Acta Crystallogr. D Biol. Crystallogr. 66, 486–501. 10.1107/S0907444910007493

Feng, S., Peng, Y., Liu, E., Ma, H., Qiao, K., Zhou, A., Liu, S., Bu, Y., 2020. Arabidopsis V-ATPase d2 Subunit Plays a Role in Plant Responses to Oxidative Stress. Genes 11, 701. 10.3390/genes11060701

Francis, M., Daniel, R., Nathalie, L.-C., 2005. New insights into the tonoplast architecture of plant vacuoles and vacuolar dynamics during osmotic stress. BMC Plant Biol. 5, 13. 10.1186/1471-2229-5-13

Goddard, T.D., Huang, C.C., Meng, E.C., Pettersen, E.F., Couch, G.S., Morris, J.H., Ferrin, T.E., 2018. UCSF ChimeraX: Meeting modern challenges in visualization and analysis. Protein Sci. Publ. Protein Soc. 27, 14–25. 10.1002/pro.3235

Guo, H., Franken, E., Deng, Y., Benlekbir, S., Singla Lezcano, G., Janssen, B., Yu, L., Ripstein, Z.A., Tan, Y.Z., Rubinstein, J.L., 2020. Electron-event representation data enable efficient cryoEM file storage with full preservation of spatial and temporal resolution. IUCrJ 7, 860–869. 10.1107/S205225252000929X

Hanitzsch, M., Schnitzer, D., Seidel, T., Golldack, D., Dietz, K.-J., 2007. Transcript level regulation of the vacuolar H+-ATPase subunit isoforms VHA-a, VHA-E and VHA-G in Arabidopsis thaliana. Mol. Membr. Biol. 24, 507–518. 10.1080/09687680701447393

Hemsley, P.A., Taylor, L., Grierson, C.S., 2008. Assaying protein palmitoylation in plants. Plant Methods 4, 2. 10.1186/1746-4811-4-2

Hong-Hermesdorf, A., Brüx, A., Grüber, A., Grüber, G., Schumacher, K., 2006. A WNK kinase binds and phosphorylates V-ATPase subunit C. FEBS Lett. 580, 932–939. 10.1016/j.febslet.2006.01.018

Jamali, K., Käll, L., Zhang, R., Brown, A., Kimanius, D., Scheres, S.H.W., 2024. Automated model building and protein identification in cryo-EM maps. Nature 628, 450–457. 10.1038/s41586-024-07215-4

Jaquinod, M., Villiers, F., Kieffer-Jaquinod, S., Hugouvieux, V., Bruley, C., Garin, J., Bourguignon, J., 2007. A proteomics dissection of Arabidopsis thaliana vacuoles isolated from cell culture. Mol. Cell. Proteomics MCP 6, 394–412. 10.1074/mcp.M600250-MCP200

Jumper, J., Evans, R., Pritzel, A., Green, T., Figurnov, M., Ronneberger, O., Tunyasuvunakool, K., Bates, R., Žídek, A., Potapenko, A., Bridgland, A., Meyer, C., Kohl, S.A.A., Ballard, A.J., Cowie, A., Romera-Paredes, B., Nikolov, S., Jain, R., Adler, J., Back, T., Petersen, S., Reiman, D., Clancy, E., Zielinski, M., Steinegger, M., Pacholska, M., Berghammer, T., Bodenstein, S., Silver, D., Vinyals, O., Senior, A.W., Kavukcuoglu, K., Kohli, P., Hassabis, D., 2021. Highly accurate protein structure prediction with AlphaFold. Nature 596, 583–589. 10.1038/s41586-021-03819-2

Kane, P.M., 1995. Disassembly and Reassembly of the Yeast Vacuolar H+-ATPase in Vivo(*). J. Biol. Chem. 270, 17025–17032. 10.1016/S0021-9258(17)46944-4

Kane, P.M., Parra, K.J., 2000. Assembly and Regulation of the Yeast Vacuolar H+-ATPase. J. Exp. Biol. 203, 81–87. 10.1242/jeb.203.1.81

Khan, Md.M., Ebrahimi, R., Oot, R.A., Wilkens, S., 2025. Interaction of yeast V-ATPase with TLDc protein Rtc5p. bioRxiv 2025.05.24.655954.

Khan, Md.M., Lee, S., Couoh-Cardel, S., Oot, R.A., Kim, H., Wilkens, S., Roh, S.-H., 2022. Oxidative stress protein Oxr1 promotes V-ATPase holoenzyme disassembly in catalytic activity-independent manner. EMBO J. 41, e109360. 10.15252/embj.2021109360

Kinouchi, K., Ichihara, A., Sano, M., Sun-Wada, G.-H., Wada, Y., Kurauchi-Mito, A., Bokuda, K., Narita, T., Oshima, Y., Sakoda, M., Tamai, Y., Sato, H., Fukuda, K., Itoh, H., 2010. The (Pro)renin Receptor/ATP6AP2 is Essential for Vacuolar H+-ATPase Assembly in Murine Cardiomyocytes. Circ. Res. 107, 30–34. 10.1161/CIRCRESAHA.110.224667

Klössel, S., Zhu, Y., Amado, L., Bisinski, D.D., Ruta, J., Liu, F., González Montoro, A., 2024. Yeast TLDc domain proteins regulate assembly state and subcellular localization of the V-ATPase. EMBO J. 43, 1870–1897. 10.1038/s44318-024-00097-2

Konrad, S.S.A., Popp, C., Stratil, T.F., Jarsch, I.K., Thallmair, V., Folgmann, J., Marín, M., Ott, T., 2014. S-acylation anchors remorin proteins to the plasma membrane but does not primarily determine their localization in membrane microdomains. New Phytol. 203, 758–769. 10.1111/nph.12867

Krebs, M., Beyhl, D., Görlich, E., Al-Rasheid, K.A.S., Marten, I., Stierhof, Y.-D., Hedrich, R., Schumacher, K., 2010. Arabidopsis V-ATPase activity at the tonoplast is required for efficient nutrient storage but not for sodium accumulation. Proc. Natl. Acad. Sci. 107, 3251–3256. 10.1073/pnas.0913035107

Krebs, M., Held, K., Binder, A., Hashimoto, K., Den Herder, G., Parniske, M., Kudla, J., Schumacher, K., 2012. FRET-based genetically encoded sensors allow high-resolution live cell imaging of Ca^2+^ dynamics. Plant J. 69, 181–192. 10.1111/j.1365-313X.2011.04780.x

Kriegel, A., Andrés, Z., Medzihradszky, A., Krüger, F., Scholl, S., Delang, S., Patir-Nebioglu, M.G., Gute, G., Yang, H., Murphy, A.S., Peer, W.A., Pfeiffer, A., Krebs, M., Lohmann, J.U., Schumacher, K., 2015. Job Sharing in the Endomembrane System: Vacuolar Acidification Requires the Combined Activity of V-ATPase and V-PPase. Plant Cell 27, 3383–3396. 10.1105/tpc.15.00733

Krüger, F., Schumacher, K., 2018. Pumping up the volume − vacuole biogenesis in Arabidopsis thaliana. Semin. Cell Dev. Biol., Redox signalling in development and regeneration 80, 106–112. 10.1016/j.semcdb.2017.07.008

Kumar, M., Carr, P., Turner, S., 2022. An atlas of Arabidopsis protein S-Acylation reveals its widespread role in plant cell organisation and function. Nat. Plants 8, 670–681. 10.1038/s41477-022-01164-4

Lampropoulos, A., Sutikovic, Z., Wenzl, C., Maegele, I., Lohmann, J.U., Forner, J., 2013. GreenGate - A Novel, Versatile, and Efficient Cloning System for Plant Transgenesis. PLOS ONE 8, e83043. 10.1371/journal.pone.0083043

Li, Y., Zeng, H., Xu, F., Yan, F., Xu, W., 2022. H+-ATPases in Plant Growth and Stress Responses. Annu. Rev. Plant Biol. 73, 495–521. 10.1146/annurev-arplant-102820-114551

Lupanga, U., Röhrich, R., Askani, J., Hilmer, S., Kiefer, C., Krebs, M., Kanazawa, T., Ueda, T., Schumacher, K., 2020. The Arabidopsis V-ATPase is localized to the TGN/EE via a seed plant-specific motif. eLife 9, e60568. 10.7554/eLife.60568

Marr, C.R., Benlekbir, S., Rubinstein, J.L., 2014. Fabrication of carbon films with ∼ 500nm holes for cryo-EM with a direct detector device. J. Struct. Biol. 185, 42–47. 10.1016/j.jsb.2013.11.002

Martinoia, E., Maeshima, M., Neuhaus, H.E., 2007. Vacuolar transporters and their essential role in plant metabolism. J. Exp. Bot. 58, 83–102. 10.1093/jxb/erl183

Maxson, M.E., Abbas, Y.M., Wu, J.Z., Plumb, J.D., Grinstein, S., Rubinstein, J.L., 2022. Detection and quantification of the vacuolar H+ATPase using the Legionella effector protein SidK. J. Cell Biol. 221, e202107174. 10.1083/jcb.202107174

Mazhab-Jafari, M.T., Rohou, A., Schmidt, C., Bueler, S.A., Benlekbir, S., Robinson, C.V., Rubinstein, J.L., 2016. Atomic model for the membrane-embedded VO motor of a eukaryotic V-ATPase. Nature 539, 118–122.

McKay, D., Lupanga, U., Uebele Perez, M., Krebs, M., Wege, S., Grabe, M., Schumacher, K., 2025. Detection of Cytosolic Ion Concentrations by the Trans-Golgi Network/Early Endosome is2 Important for Salt Tolerance. bioRxiv 13.670069.

Mencia, R., Céccoli, G., Fabro, G., Torti, P., Colombatti, F., Ludwig-Müller, J., Alvarez, M.E., Welchen, E., 2020. OXR2 Increases Plant Defense against a Hemibiotrophic Pathogen via the Salicylic Acid Pathway. Plant Physiol. 184, 1112–1127. 10.1104/pp.19.01351

Merkulova, M., Păunescu, T.G., Azroyan, A., Marshansky, V., Breton, S., Brown, D., 2015. Mapping the H(+) (V)-ATPase interactome: identification of proteins involved in trafficking, folding, assembly and phosphorylation. Sci. Rep. 5, 14827. 10.1038/srep14827

Nguyen, J.T., Ray, C., Fox, A.L., Mendonça, D.B., Kim, J.K., Krebsbach, P.H., 2018. Mammalian EAK-7 activates alternative mTOR signaling to regulate cell proliferation and migration. Sci. Adv. 4, eaao5838. 10.1126/sciadv.aao5838

Oot, R.A., Wilkens, S., 2024. Human V-ATPase function is positively and negatively regulated by TLDc proteins. Struct. Lond. Engl. 1993 32, 989–1000.e6. 10.1016/j.str.2024.03.009

Park, J., Song, W.-Y., Ko, D., Eom, Y., Hansen, T.H., Schiller, M., Lee, T.G., Martinoia, E., Lee, Y., 2012. The phytochelatin transporters AtABCC1 and AtABCC2 mediate tolerance to cadmium and mercury. Plant J. 69, 278–288. 10.1111/j.1365-313X.2011.04789.x

Parra, K., Keenan, K., Kane, P., 2000. The H subunit (VMA13p) of the yeast V-ATPase inhibits the ATPase activity of cytosolic V1 complexes. J. Biol. Chem. 275, 21761–21767. 10.1074/jbc.M002305200

Punjani, A., Rubinstein, J.L., Fleet, D.J., Brubaker, M.A., 2017. cryoSPARC: algorithms for rapid unsupervised cryo-EM structure determination. Nat. Methods 14, 290–296. 10.1038/nmeth.4169

Qi, J., Forgac, M., 2008. Function and subunit interactions of the N-terminal domain of subunit a (Vph1p) of the yeast V-ATPase. J Biol Chem 283, 19274–19282. 10.1074/jbc.M802442200

Rappsilber, J., Ishihama, Y., Mann, M., 2003. Stop and go extraction tips for matrix-assisted laser desorption/ionization, nanoelectrospray, and LC/MS sample pretreatment in proteomics. Anal. Chem. 75, 663–670. 10.1021/ac026117i

Roh, S.-H., Stam, N.J., Hryc, C.F., Couoh-Cardel, S., Pintilie, G., Chiu, W., Wilkens, S., 2018. The 3.5-Å CryoEM Structure of Nanodisc-Reconstituted Yeast Vacuolar ATPase V o Proton Channel. Mol. Cell 69, 993–1004. 10.1016/j.molcel.2018.02.006

Rubinstein, J.L., Brubaker, M.A., 2015. Alignment of cryo-EM movies of individual particles by optimization of image translations. J. Struct. Biol. 192, 1–11. 10.1016/j.jsb.2015.08.007

Schindelin, J., Arganda-Carreras, I., Frise, E., Kaynig, V., Longair, M., Pietzsch, T., Preibisch, S., Rueden, C., Saalfeld, S., Schmid, B., Tinevez, J.-Y., White, D.J., Hartenstein, V., Eliceiri, K., Tomancak, P., Cardona, A., 2012. Fiji: an open-source platform for biological-image analysis. Nat. Methods 9, 676–682. 10.1038/nmeth.2019

Schulze, W.X., Schneider, T., Starck, S., Martinoia, E., Trentmann, O., 2012. Cold acclimation induces changes in Arabidopsis tonoplast protein abundance and activity and alters phosphorylation of tonoplast monosaccharide transporters. Plant J. Cell Mol. Biol. 69, 529–541. 10.1111/j.1365-313X.2011.04812.x

Schumacher, K., Krebs, M., 2010. The V-ATPase: small cargo, large effects. Curr. Opin. Plant Biol. 13, 724–730. 10.1016/j.pbi.2010.07.003

Schwanhäusser, B., Busse, D., Li, N., Dittmar, G., Schuchhardt, J., Wolf, J., Chen, W., Selbach, M., 2011. Global quantification of mammalian gene expression control. Nature 473, 337–342. 10.1038/nature10098

Sorek, N., Segev, O., Gutman, O., Bar, E., Richter, S., Poraty, L., Hirsch, J.A., Henis, Y.I., Lewinsohn, E., Jürgens, G., Yalovsky, S., 2010. An S-acylation switch of conserved G domain cysteines is required for polarity signaling by ROP GTPases. Curr. Biol. CB 20, 914–920. 10.1016/j.cub.2010.03.057

Strazzer, P., Spelt, C.E., Li, S., Bliek, M., Federici, C.T., Roose, M.L., Koes, R., Quattrocchio, F.M., 2019. Hyperacidification of Citrus fruits by a vacuolar proton-pumping P-ATPase complex. Nat. Commun. 10, 744. 10.1038/s41467-019-08516-3

Strompen, G., Dettmer, J., Stierhof, Y.-D., Schumacher, K., Jürgens, G., Mayer, U., 2005. Arabidopsis vacuolar H+-ATPase subunit E isoform 1 is required for Golgi organization and vacuole function in embryogenesis. Plant J. 41, 125–132. 10.1111/j.1365-313X.2004.02283.x

Sun, N., Xu, X., Zhu, Z., Zhou, X., Liu, Y., Li, D., Cao, F., Wang, L., Zhang, H., 2025. Tonoplast sugar transporter ZmTST1 positively regulates plant growth, salt and drought tolerance. Plant Physiol. Biochem. 229, 110380. 10.1016/j.plaphy.2025.110380

Sze, H., Schumacher, K., Müller, M.L., Padmanaban, S., Taiz, L., 2002. A simple nomenclature for a complex proton pump: VHA genes encode the vacuolar H+-ATPase. Trends Plant Sci. 7, 157–161. 10.1016/S1360-1385(02)02240-9

Tan, Y.Z., Abbas, Y.M., Wu, J.Z., Wu, D., Keon, K.A., Hesketh, G.G., Bueler, S.A., Gingras, A.-C., Robinson, C.V., Grinstein, S., Rubinstein, J.L., 2022a. CryoEM of endogenous mammalian V-ATPase interacting with the TLDc protein mEAK-7. Life Sci. Alliance 5. 10.26508/lsa.202201527

Tan, Y.Z., Keon, K.A., Abdelaziz, R., Imming, P., Schulze, W., Schumacher, K., Rubinstein, J.L., 2022b. Structure of V-ATPase from citrus fruit. Structure 30, 1403–1410.e4. 10.1016/j.str.2022.07.006

Vasanthakumar, T., Keon, K.A., Bueler, S.A., Jaskolka, M.C., Rubinstein, J.L., 2022. Coordinated conformational changes in the V1 complex during V-ATPase reversible dissociation. Nat. Struct. Mol. Biol. 29, 430–439. 10.1038/s41594-022-00757-z

Viotti, C., Bubeck, J., Stierhof, Y.-D., Krebs, M., Langhans, M., Van Den Berg, W., Van Dongen, W., Richter, S., Geldner, N., Takano, J., Jürgens, G., De Vries, S.C., Robinson, D.G., Schumacher, K., 2010. Endocytic and Secretory Traffic in *Arabidopsis* Merge in the Trans-Golgi Network/Early Endosome, an Independent and Highly Dynamic Organelle. Plant Cell 22, 1344–1357. 10.1105/tpc.109.072637

Volkert, M.R., Elliott, N.A., Housman, D.E., 2000. Functional genomics reveals a family of eukaryotic oxidation protection genes. Proc. Natl. Acad. Sci. 97, 14530–14535. 10.1073/pnas.260495897

Wang, H., Bueler, S.A., Rubinstein, J.L., 2023. Structural basis of V-ATPase VO region assembly by Vma12p, 21p, and 22p. Proc. Natl. Acad. Sci. U. S. A. 120, e2217181120. 10.1073/pnas.2217181120

Wang, H., Rubinstein, J.L., 2023. CryoEM of V-ATPases: Assembly, disassembly, and inhibition. Curr. Opin. Struct. Biol. 80, 102592. 10.1016/j.sbi.2023.102592

Wang, L., Wu, D., Robinson, C.V., Fu, T.-M., 2022. Identification of mEAK-7 as a human V-ATPase regulator via cryo-EM data mining. Proc. Natl. Acad. Sci. 119, e2203742119. 10.1073/pnas.2203742119

Wang, R., Long, T., Hassan, A., Wang, J., Sun, Y., Xie, X.S., Li, X., 2020. Cryo-EM structures of intact V-ATPase from bovine brain. Nat. Commun. 11, 1–9. 10.1038/s41467-020-17762-9

Wang, R., Qin, Y., Xie, X.-S., Li, X., 2022. Molecular basis of mEAK7-mediated human V-ATPase regulation. Nat. Commun. 13, 3272. 10.1038/s41467-022-30899-z

Yamauchi, S., Fusada, N., Hayashi, H., Utsumi, T., Uozumi, N., Endo, Y., Tozawa, Y., 2010. The consensus motif for N-myristoylation of plant proteins in a wheat germ cell-free translation system. FEBS J. 277, 3596–3607. 10.1111/j.1742-4658.2010.07768.x

Zauber, H., Schüler, V., Schulze, W.X., 2013. Systematic Evaluation of Reference Protein Normalization in Proteomic Experiments. Front. Plant Sci. 4. 10.3389/fpls.2013.00025

Zhang, X., Henriques, R., Lin, S.-S., Niu, Q.-W., Chua, N.-H., 2006. Agrobacterium-mediated transformation of Arabidopsis thaliana using the floral dip method. Nat. Protoc. 1, 641–646. 10.1038/nprot.2006.97

Zhao, J., Benlekbir, S., Rubinstein, J.L., 2015. Electron cryomicroscopy observation of rotational states in a eukaryotic V-ATPase. Nature 521, 241–245. 10.1038/nature14365

